# PHR1 and PHL1 mediate rapid high-light responses and acclimation to high photosynthetic activity

**DOI:** 10.1101/2023.03.11.532192

**Authors:** Lukas Ackermann, Agnes Fekete, Laura Schröder, Thomas Nägele, Alina J. Hieber, Monika Müller, Martin J. Mueller, Maria Klecker

## Abstract

- Rapid acclimation towards high light is vital for the prevention of phototoxic stress to plant tissues. Photosynthesis transiently consumes inorganic phosphate (P_i_). We therefore asked whether changes in intracellular P_i_ pools can act as a signal for reprogramming gene expression upon increased light intensity.
- The function of triose phosphate utilization for intracellular P_i_ homeostasis was deciphered by investigation of mutants defective in both photoassimilate partitioning and low-P_i_ signaling. Next, we determined subcellular P_i_ levels and transcript accumulation upon short-term high light. Physiological consequences were examined by analyses of ATP, carbohydrates, and an untargeted lipid profiling approach.
- The capacity for triose phosphate utilization clearly affected systemic P_i_ signaling. High-light treatment caused a rapid depletion specifically of chloroplast P_i_ levels paralleled by induction of transcripts dependent on PHOSPHATE STARVATION RESPONSE 1 (PHR1) and PHR1-LIKE 1. Among the high-light induced targets of PHR1, *SRG3/GDPD1* is involved in phospholipid catabolism. Lipid profiling revealed differences between WT and *srg3* mutant plants upon high light, including changes in linolenic acid and photoprotective zeaxanthin.
- We conclude that photosynthetic activity regulates the low-P_i_ response machinery in the nucleus to implement high-light acclimation. This facilitates the liberation of cellular P as well as the maintenance of membrane lipid integrity.

## Introduction

Light energy is provided to photoautotrophic organisms at variable intensities that can fall below or exceed the requirements of the photosynthetic machinery at unpredictable times during the day period. This necessitates an enzymatic machinery capable of coping with rapid changes in metabolic flux, as well as the containment of overexcitation-induced hazards, such as lipid peroxidation (Havaux & Niyogi, 1999). Both challenges are met not only by post-transcriptional regulation of enzyme activity (reviewed in König et al. (2012), Matiolli et al. (2022)), but also by pronounced changes in mRNA abundance that occur within 30 minutes upon an increase in irradiance (Vogel *et al*., 2014; Suzuki *et al*., 2015; Huang *et al*., 2019). The latter mechanism depends on fast signal transmission between the chloroplast and the nuclear transcription machinery. Such communication is mediated by ‘primary’ retrograde signals (Dietz, 2015), including redox cues and reactive oxygen species which are produced during photosynthetic reactions (Bechtold *et al*., 2008; Dietz *et al*., 2016). Additionally, increased electron transport activity and light-induced Calvin-Benson cycle activation promote the accumulation of primary products of carbon fixation (Dietz & Heber, 1984, 1986; Borghi *et al*., 2019) which may themselves act as signaling molecules (Häusler *et al*., 2014; Moore *et al*., 2014). Hence, induction of high-light specific gene expression partially depends on the triose phosphate/phosphate translocator (TPT) that mediates the exchange of triose phosphates and 3-phosphoglycerate (3-PGA) with inorganic phosphate (P_i_) at the chloroplast envelope (Schneider *et al*., 2002; Vogel *et al*., 2014; Weise *et al*., 2019). A central function of the TPT in light acclimation and retrograde signaling was corroborated by the severe high-light dependent phenotypes of double mutants defective in *TPT* and *ADG1* (for *ADP GLUCOSE PYROPHOSPHORYLASE 1*), encoding a small subunit of the ADP-glucose pyrophosphorylase complex (AGPase) which is required for the biosynthesis of transitory starch (Schmitz *et al*., 2012). Phenotypes of *adg1-1 tpt* double mutants include growth retardation, low diurnal changes in carbohydrate levels, and high chlorophyll fluorescence. These observations point towards an important role of photosynthate partitioning for retrograde signaling, however the identity of the primary sensors as well as signal transduction mechanisms are still unclear.

Phosphorus makes up about 2-30 permille of the plant dry mass (Kumar *et al*., 2019), as it is incorporated into a large subset of biomolecules. Systemic signaling of P_i_ availability and P_i_ distribution within the plant body mainly depend on a conserved gene family encoding PHOSPHATE STARVATION RESPONSE 1 (PHR1) and at least 14 PHR1 LIKE (PHL) transcription factors of the GARP coiled-coiled type (Rubio *et al*., 2001; Zhou *et al*., 2008; Bustos *et al*., 2010; Thibaud *et al*., 2010; Sun *et al*., 2016; Safi *et al*., 2017). Simultaneous loss of *PHR1* and *PHL1* function affects the expression of roughly 70 % of the P_i_ starvation responsive genes in the shoots of Arabidopsis (*Arabidopsis thaliana)* (Bustos *et al*., 2010). A decrease in cellular P_i_ concentration impairs production of the inositol pyrophosphate species InsP8 in the cytosol, relieving PHR1/PHLs of suppression by SPX (for SYG1/Pho81/XPR1) domain-containing proteins, thus linking transcription factor activity to P_i_ availability (Puga *et al*., 2014; Dong *et al*., 2019; Zhu *et al*., 2019; Ried *et al*., 2021). The transcriptional response to P_i_ scarcity leads to scavenging of external and internal P_i_ sources through phosphatase induction (Morcuende *et al*., 2007) and altered lipid metabolism (Misson *et al*., 2005; Pant *et al*., 2015; Yang *et al*., 2025), anthocyanin biosynthesis (Nilsson *et al*., 2012; Liu *et al*., 2022; Li *et al*., 2023), and protection from photodamage (Nilsson *et al*., 2012), to name a selection. The largest pool of cellular P reserves resides in the vacuole, and it has been shown in rice (*Oryza sativa*), that vacuolar P_i_ efflux into the cytosol is mediated by two proteins of the glycerol 3-phosphate transporter family which are transcriptionally induced by PHR proteins (Xu *et al*., 2019). Three homologs have been identified in Arabidopsis and are termed VPE1-3 (Ramaiah *et al*., 2011; Xu *et al*., 2019), but their contribution to P_i_ homeostasis has not been addressed so far in dicotyledonous plants. PHR1/PHLs are characterized as transcriptional activators (Nilsson *et al*., 2007; Bustos *et al*., 2010). Accordingly, direct targets of PHR1 (Bustos *et al*., 2010; Castrillo *et al*., 2017) are strongly upregulated upon P_i_ starvation and include genes such as *PHOSPHATE STARVATION-INDUCED GENE 2* (*PS2*; AT1G73010) (Hanchi *et al*., 2018), *MONOGALACTOSYLDIACYLGLYCEROL SYNTHASE 3* (*MGD3*; AT2G11810) (Kobayashi *et al*., 2004), and *GLYCEROPHOSPHODIESTER PHOSPHODIESTERASE 1/SENESCENCE-RELATED GENE 3* (*GDPD1*/*SRG3*; AT3G02040) (Cheng *et al*., 2011). The latter encodes a plastid-localized enzyme with broad-spectrum glycerophosphodiester phosphodiesterase activity releasing primary alcohols and glycerol 3-phosphate from deacylated phospholipids (Cheng *et al*., 2011). SRG3 is supposed to mainly act in the catalysis of phosphatidylcholine to provide glycerol-3-phosphate for DGDG biogenesis, for glycolytic utilization, or liberation of P_i_ by subsequent acid phosphatase activity (Cheng *et al*., 2011). A function in phospholipid homeostasis was recently corroborated by Wojciechowska et al. (2024) who proposed that the transcriptional repressor PROTODERMAL FACTOR2 directly links phospholipid availability to the repression of both *SRG3* and *SPX1*. The *SRG3* gene was shown to be important for seedling fitness and maintenance of cellular P_i_ levels under P_i_ starvation conditions (Cheng *et al*., 2011), but how exactly SRG3 contributes to nutrient homeostasis is not understood.

P_i_ is unique among the macronutrients in that it is the only root-supplied element which is directly required for photosynthesis (Heldt & Rapley, 1970; Dietz & Heber, 1984). It has been shown experimentally that under high-CO_2_ and high-light conditions, photosynthesis can indeed become limited by P_i_ supply to the chloroplast, both under P_i_ depleted and nutrient-rich conditions (Walker & Osmond, 1986; Dietz & Foyer, 1986; Sivak & Walker, 1986). However, whether rapid increase in photosynthetic activity directly affects P_i_ homeostasis of the cell, is still unclear. In this study, we show that increments in light intensity rapidly deplete chloroplast P_i_ reserves *in planta*. Strikingly, although other cellular P_i_ pools are not affected by short-term high light, this is accompanied by a nuclear transcriptional response characteristic of P_i_ depletion which depends on *PHR1* and *PHL1*. Transcript analysis of the *adg1 tpt-2* mutant links PHR(-like) activation to triose phosphate allocation. Processes regulated by PHR1/PHL1 upon high light contribute to photosynthetic acclimation since *phr1 phl1* mutants are sensitive to high-light growth conditions, and showed altered carbohydrate and anthocyanin accumulation compared to the wildtype (WT) when the light intensity was increased. Moreover, analyses of the starch biosynthesis-deficient *adg1 phr1 phl1* triple mutant revealed an important function of P_i_ signaling when triose phosphate utilization is constrained. Induction of genes involved in lipid metabolism such as *SRG3* by PHR1/PHL1 furthermore contributes to fatty acid homeostasis and membrane photoprotection, as revealed by lipid profiling of the *srg3* mutant upon high light. Together, we propose that PHR1/PHL1 mediate cellular P_i_ release and rapid photosynthetic acclimation upon increases in light intensity, probably activated by direct sensing of chloroplast P_i_ levels upon triose phosphate overproduction.

## Results

### Triose phosphate utilization affects P_i_ signaling

Photoassimilate allocation and utilization in the chloroplast and cytosol are supposed to limit the rate of P_i_ recycling from photosynthetic products (McClain & Sharkey, 2019). In line with this assumption, P_i_ deprivation has been shown to stimulate starch accumulation in a *PHR1*-dependent manner, which involves AGPase activity (Nilsson *et al*., 2007). To address whether partitioning of phosphorylated assimilates contributes to P_i_ homeostasis of the cell, we examined the phenotypes of mutants defective in starch biosynthesis (*adg1-1*) and chloroplastic triose phosphate export (*tpt-2*) during P_i_ starvation in the presence of 14.61 mM (0.5 %) sucrose. Starch accumulation in response to P_i_ depletion was abolished in both the *phr1-3 phl1* double mutant and the *adg1-1* mutant (Figure 1A), confirming that PHR(-like) signaling as well as the small AGPase subunit ADG1 are required for increased starch production under P_i_ starvation. Concomitantly, P_i_-depleted *adg1-1* mutants accumulated high levels of monosaccharides (glucose and fructose, Figure 1A). Loss of TPT function caused significantly higher levels of starch and glucose under P_i_ deprivation as compared to the WT, which resumed a trend that was also observed under control conditions and likely reflects increased day-time starch turnover in the *tpt-2* mutant (Schneider *et al*., 2002). Interestingly, over-accumulation of carbohydrates by *adg1-1* and *tpt-2* mutants under P_i_ deprivation was ceased in the double mutant *adg1 tpt-2*, indicating that retrograde signaling of carbohydrate status is involved in this process. Likewise, *adg1 tpt-2* double mutants failed to accumulate anthocyanins upon P_i_ deprivation (Supporting Information Fig. S1A, B).

**Figure 1.**
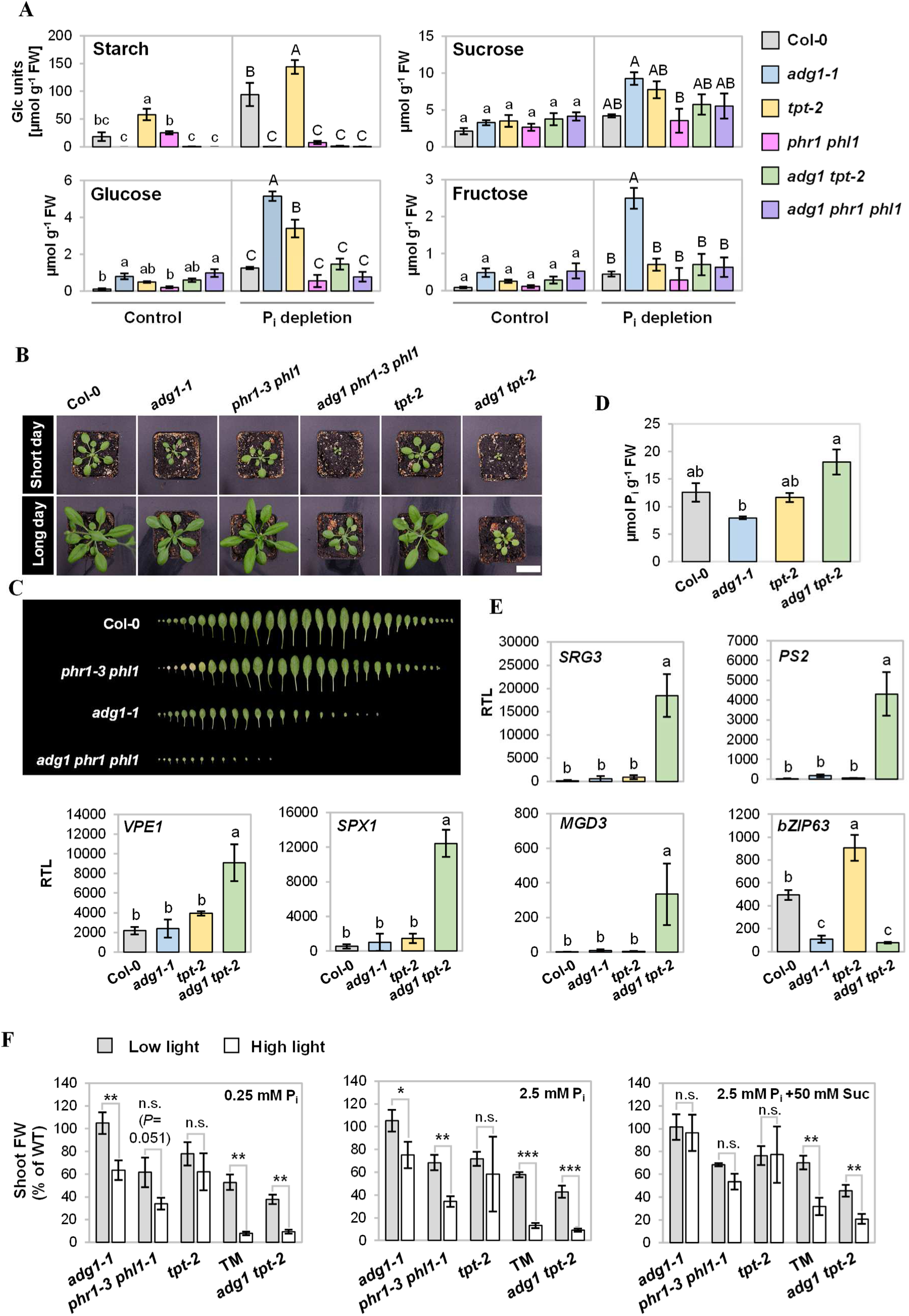
Mutants with defects in photoassimilate partitioning show altered P_i_ responses. **A,** Contents of glucose, fructose, sucrose, and starch in shoots upon P_i_ depletion relative to fresh weight (**FW**). Seedlings of WT (Col-0), *adg1-1*, *tpt-2*, *phr1-3 phl1*, *adg1 tpt-2* and *adg1 phr1-3 phl1* genotype were grown for 10 days on rich medium containing 0.5 % sucrose before transfer to media with 0.5 % sucrose and either 2.5 mM (**Control**) or 0 mM (**P_i_ depletion**) KH_2_PO_4_ added. Growth was continued for 7 days. Pools of seedling shoots were harvested 10.25 h after onset of the 16-h photoperiod. Means ± standard deviations; *n* = 3 independent experiments. Letter code indicates statistical differences at *P* < 0.05. One-way ANOVA with Tukey HSD follow-up test and Bonferroni alpha correction for contrasts. **B, C,** Phenotype of the *adg1 phr1 phl1* triple mutant. Plants were grown at 23°C and a light intensity of 100±10 µmol m^-2^ s^-1^. **B,** Rosette habitus of plants grown under short-day (8 h light period) and long-day (16 h light period) conditions. Pictures were taken of representative plants after 27 (short day) and 25 (long day) days of growth. Bar, 2.5 cm. **C,** Leaves of 42 day-old plants grown under an 8-h light regime were aligned according to age (starting with the cotyledons on the left hand side). Background was manually removed for better visualization. **D, E,** Constitutive P_i_-starvation transcription in rosette leaves of *adg1 tpt-2*. Plants were grown on soil at a photon flux density of 90±10 µmol m^-2^ s^-1^ and sampled 135 min after onset of the 8-h photoperiod after 5 weeks of growth. **D,** P_i_ contents of rosette leaves. Means ± standard deviations are depicted. *n* = 3 independent experiments. Kruskal-Wallis test with DUNN‘s follow-up test and DUNN/Sidak alpha correction for contrasts; *P* < 0.01. **E,** qRT-PCR analysis of P_i_ starvation responsive genes in rosette leaves of WT, *adg1-1*, *tpt-2*, and *adg1 tpt-2*. Transcript levels are 1000·2^-ΔCT^ relative to *PP2A*. Bars show means ± standard deviations. *n* = 3 independent experiments. One-way ANOVA with Tukey HSD follow-up test and Bonferroni alpha correction for contrasts; *P* < 0.01. **F**, High-light sensitivity of *adg1-1*, *phr1-3 phl1* and derived mutants. Seedlings were grown for two weeks in a 16/8 h (light/dark) cycle under low light (60±5 µmol m^-2^ s^-1^) or high light (350±30 µmol m^-2^ s^-1^) on agar-solidified media containing 0.25 mM P_i_, 2.5 mM P_i_, or 2.5 mM P_i_ +50 mM sucrose. Shoot FW were calculated as percent of mean shoot FW of WT seedlings grown under the respective conditions. Bars represent means ± standard deviations; *n =* 3 independent experiments comprising means of each 4-8 seedling shoots. Statistical analyses were performed using Student’s *t* test with 2-tailed distribution, unpaired with unequal variance. Significant differences between data are indicated as \*\*\**P* < 0.001, \*\**P* < 0.01, \**P* < 0.05, n.s., not significant.

In order to test if nuclear gene regulation by PHR1/PHL1 is important for the adjustment of P homeostasis when P_i_ recycling from photosynthetic intermediates is impaired, we generated the triple mutant *adg1-1 phr1-3 phl1* (hereafter referred to as *adg1 phr1 phl1*). This mutant exhibited a pronounced growth defect which was the result of reduced leaf size and number (Figure 1B, C). Next, *adg1 phr1 phl1* plants flowered later than the parental lines (Supporting Information Fig. S1C). These findings indicate that starch production functions synergistically with PHR1/PHL1 in maintaining cellular P_i_ and/or carbohydrate homeostasis.

We noted that growth retardation of *adg1 phr1 phl1* resembled the phenotype of the *adg1 tpt-2* mutant (Figure 1B). To test the hypothesis that *adg1 tpt-2* might likewise be affected in P_i_ homeostasis, we measured P_i_ contents of aboveground tissues and determined P_i_ starvation gene expression in vegetative plants grown on soil. Surprisingly, we found that *adg1 tpt-2* double mutants over-accumulated P_i_ compared to the *adg1-1* single mutant (Figure 1D). Counter-intuitively, quantitative RT-PCR (qRT-PCR) analysis revealed that transcripts of P_i_ starvation marker genes including *SRG3*, *PS2*, *VPE1* (AT3G47420), *SPX1*, and *MGD3* (Supporting Information Fig. S2) were all constitutively up-regulated in *adg1 tpt-2* (Figure 1E). Additionally, transcript abundance of the circadian regulator *BASIC LEUCINE ZIPPER 63* (*bZIP63*) (Baena-González *et al*., 2007; Frank *et al*., 2018), which is repressed by P_i_ depletion in the WT (Supporting Information Fig. S2), was also reduced in rosettes of *adg1 tpt-2* as well as *adg1-1* when grown on soil (Figure 1E). Interestingly, *tpt-2* on the contrary showed slightly higher expression levels of *bZIP63* than WT plants. Thus, P_i_ signaling pathways are substantially affected when triose phosphate utilization and allocation in the chloroplast are compromised.

It was reported that the *adg1 tpt-2* mutant is sensitive towards higher light intensities, which could be overcome by exogenously supplied sucrose (Schmitz *et al*., 2012; Heinrichs *et al*., 2012). Given the phenotypic similarities between *adg1 tpt-2* and *adg1 phr1 phl1,* we asked if the growth defect of *adg1 phr1 phl1* was also dependent on light intensity. In fact, we observed high-light dependent growth restriction of *adg1 phr1 phl1* seedlings that was comparable to *adg1 tpt-2*, both under low and high P_i_ conditions (Figure 1F; Supporting Information Fig. S3). As for *adg1 tpt-2*, the difference in growth performance between low-and high-light conditions was alleviated by providing sucrose to *adg1 phr1 phl1* mutants (Figure 1F), a phenomenon that was also observed for root growth of the mutant seedlings (Supporting Information Fig. S3). Remarkably, high-light dependent growth restriction was also seen for both the *adg1-1* and *phr1-3 phl1* mutants, although to a lower extend than in the cases of *adg1 tpt-2* and *adg1 phr1 phl1* (Figure 1F). Again, the difference in growth performance between low and high light was attenuated by exogenous sucrose, indicating that carbohydrate homeostasis is disturbed in *adg1-1*, *phr1-3 phl1*, and the triple mutant when exposed to high light.

We concluded from these experiments that triose phosphate allocation has pronounced effects on the nuclear P_i_ starvation response. Vice versa, P_i_ signaling is required for acclimation to triose phosphate oversupply in high light or when starch production is genetically disabled.

### High light elicits P_i_ starvation gene expression in the WT

The results of our reverse genetic analyses indicated that P_i_ homeostasis is affected by photoassimilate availability and utilization. This raised the question whether P_i_ signaling was also modulated in WT plants under non-starved conditions by naturally occurring changes in the photosynthetic rate such as upon a shift in light intensity. In order to address this, we interrogated published transcriptomic data collected from seedling shoots under P_i_ starvation (Bustos *et al*., 2010) and high-light treatments under nutrient-rich conditions (Huang *et al*., 2019). We found that out of 379 transcripts upregulated (log_2_ fold change >2) by 30 min of high light, 75 were also upregulated by P_i_ starvation (Figure 2A). For longer high-light treatments of 24 hours, 131 out of 564 transcripts were also P_i_-responsive (Figure 2A). In both cases, the overlap was found to be significant at *P* < 0.0001 in hypergeometric testing using Fisher’s exact test. In order to estimate whether the observed overlap between high-light and low-P_i_ induced transcripts might be due to generally higher nutrient requirements under high-light conditions, we next included a set of nitrogen-responsive transcripts in the comparison (Krapp *et al*., 2011). We detected a significant overlap of 33 transcripts that are differentially expressed in the shoots under low nitrogen conditions with transcripts upregulated by a high-light treatment of 24 hours duration (Figure 2A). In contrast, no significant enrichment of nitrogen-responsive transcripts among high-light induced transcripts was observed for 30-min high-light treated plants (10 out of 379). Thus, short-term high-light treatment specifically triggered low-P_i_ responses, while long-term high-light stress seems to elevate general nutrient demand of the plant.

**Figure 2.**
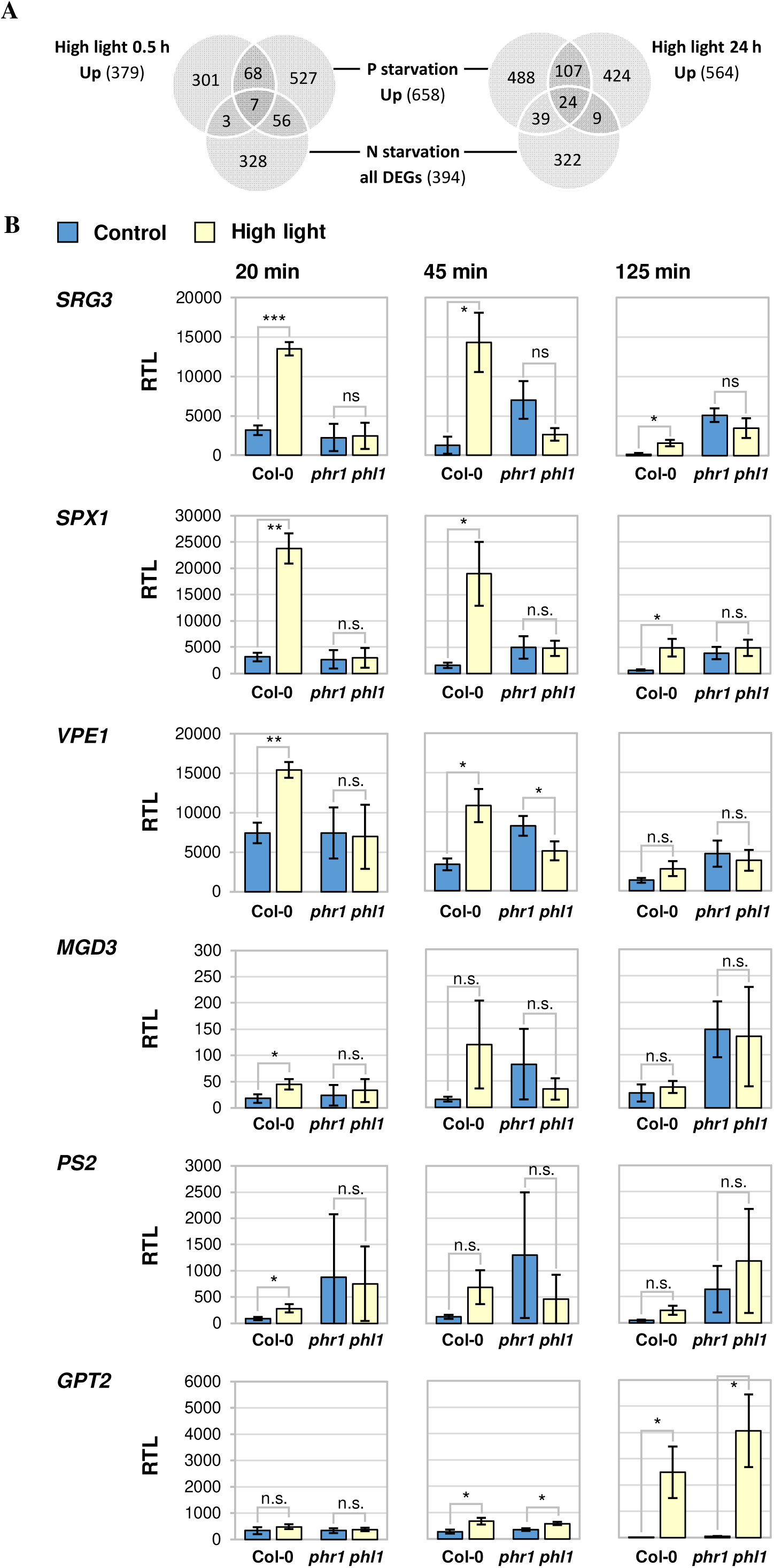
High light triggers P_i_-depletion responsive gene expression. **A**, Overlap of differential gene expression in shoots upon high light, P_i_ starvation, and nitrogen starvation. Venn diagrams show numbers of transcripts upregulated by high light or P_i_ starvation, and of genes differentially expressed under N starvation. Transcript data are taken from Huang et al. (2019) (high light), Bustos et al. (2010) (P_i_ depletion), and Krapp et al. (2011) (N depletion). **B**, qRT-PCR analyses of selected transcripts in rosette leaves from WT (Col-0) and *phr1-1 phl1* mutants. Plants were grown on soil at 23°C for 5 weeks in an 8-h light regime at 70±5 µmol m^-2^ s^-1^ before light intensity was set to 450±30 µmol m^-2^ s^-1^ (High light, yellow bars) starting 1 h after onset of the photoperiod on day 36. The control group was kept at 70±5 µmol m^-2^ s^-1^ (blue bars). Each 2-3 fully developed leaves (leaves no. 10-13) were harvested per sample at the indicated treatment times. Transcript levels were calculated relative to *PP2A* as 1000·2^-ΔCT^. Bars represent means ± standard deviations, *n* = 3 independent experiments. Statistical analyses were performed using Student’s *t* test with 2-tailed distribution, unpaired with unequal variance. Significant differences are indicated as \*\*\**P* <0.001, \*\**P* <0.01, \**P* <0.05, n.s., not significant.

We next asked whether the induction of P_i_-starvation responsive genes by short-term high light required the activity of PHR1(-like) transcription factors and thus was truly caused by low-P_i_ signaling pathways. To this end, we grew WT and *phr1-1 phl1* mutant plants under nutrient-rich conditions on soil in an 8/16 hours (light/dark) photoregime. After 5 weeks, the photon flux density was increased to 450±30 µmol m^-2^ s^-1^ (high light) at 1 hour after commencement of the photoperiod, while the control group was kept under growth light conditions of 70±5 µmol m^-2^ s^-1^. We conducted qRT-PCR analysis of samples harvested after 20, 45, and 125 min of treatment and detected pronounced upregulation of *SRG3*, *SPX1*, and *VPE1* transcripts in WT plants already after 20 min of exposure to high light compared to control plants, as well as a slight increase in *MGD3* and *PS2* transcript abundance (Figure 2B). In all five cases, the upregulation of gene expression was attenuated within 125 min in high light, confirming that response to P_i_ shortage constitutes an early consequence of increased irradiance. Crucially, P_i_-starvation marker transcripts were not responsive to light conditions in *phr1-1 phl1* mutant plants (Figure 2B), suggesting that PHR1/PHL1 are required for high-light mediated regulation of these genes. Instead, we observed a tendency towards constitutively higher transcript levels of most tested genes in the *phr1-1 phl1* mutants under control conditions, particularly within the later timepoints of the analysis. This is likely caused by activation of the remaining PHL genes in this mutant exhibiting chronically low P_i_ levels.

*PHR1* expression itself was reported to depend on light (Liu *et al*., 2017). Importantly however, increasing the light intensity to 450±30 µmol m^-2^ s^-1^ did not affect *PHR1* transcript levels under our experimental conditions (Supporting Information Fig. S4), consistent with the accumulation of *PHR1* transcript being only responsive to changes in the very low fluence range (Liu *et al*., 2017).

*GLUCOSE-6-PHOSPHATE/PHOSPHATE TRANSLOCATOR 2* (*GPT2*) is a light responsive gene (Athanasiou *et al*., 2010) which was also described to be induced by P_i_ starvation (Morcuende *et al*., 2007). Furthermore, high-light induced expression of *GPT2* was previously described to depend on *TPT* function (Kunz *et al*., 2010; Weise *et al*., 2019), and we noted that the promotor sequence of *GPT2* contains a PHR1 binding motif (GTATATTC) close to the transcriptional start site. However, in our high-light experiment, *GPT2* was upregulated in both WT and *phr1-1 phl1* after 45 minutes, and transcript levels continued to rise upon stress during the course of the analysis (Figure 2B). This showed that PHR1/PHL1 are not required for high-light mediated induction of *GPT2* expression. Moreover, we were not able to detect an effect of the *gpt2-1* allele (Niewiadomski *et al*., 2005) on carbohydrate accumulation under P_i_ depletion (Supporting Information Fig. S5), arguing against a major role of this gene for sugar homeostasis under P_i_ starvation. Thus, *GPT2* induction under high light is independent of PHR1(-like) signaling, and *phr1 phl1* mutants are not generally impaired in the transcriptional response to high light. Nevertheless, high-light treatment at early timepoints specifically triggers low-P_i_ signaling resulting in the upregulation of transcripts such as *SRG3*, *SPX1*, and *VPE1*.

### Short-term high light affects chloroplast P_i_ levels and sucrose accumulation

Upregulation of P_i_ starvation marker genes upon short-term high light suggested the existence of a light-induced signal resulting in the activation of PHR1(-like) transcription factors. Since PHR1(-like) activity is primarily regulated by the P_i_ status of the cell, we asked whether cellular P_i_ reserves were affected by high light, thus relieving PHR(-like) transcription factors of SPX-mediated inhibition. To address this, we performed non-aqueous fractionation of plant rosettes subjected to high light for 20 min or kept under control conditions, and determined subcellular concentrations of free P_i_. Under control conditions, WT and *phr1-1 phl1* mutant plants clearly differed in vacuolar P_i_ reserves (14 % of WT level in *phr1-1 phl1*), chloroplast P_i_ content (50 %), as well as the cytosolic and nuclear P_i_ fraction (63 %) (Figure 3A). Strikingly, high-light treatment caused a decline in chloroplast P_i_ levels by 54 % in the WT (Figure 3A), decreasing to 0.4 µmol P_i_ g^-1^ dry weight which corresponds to a concentration of about 0.12 mM P_i_ (Santarius & Heber, 1965). Consequently, P_i_ levels of high-light treated chloroplasts in the WT approximated the P_i_ levels measured in chloroplasts of *phr1-1 phl1* under control conditions. In contrast, no light-dependent effect was seen for P_i_ levels in the double mutant. Importantly, P_i_ concentrations of the cytosolic/nuclear and vacuolar fractions were not significantly altered by high light in either WT or *phr1-1 phl1* mutant plants (Figure 3A). Hence, elevated light rapidly induced P_i_ depletion in the WT, however the effect remained restricted to the chloroplast, and therefore does not qualify as a direct cause of PHR1/PHL1 activation upon high light.

**Figure 3.**
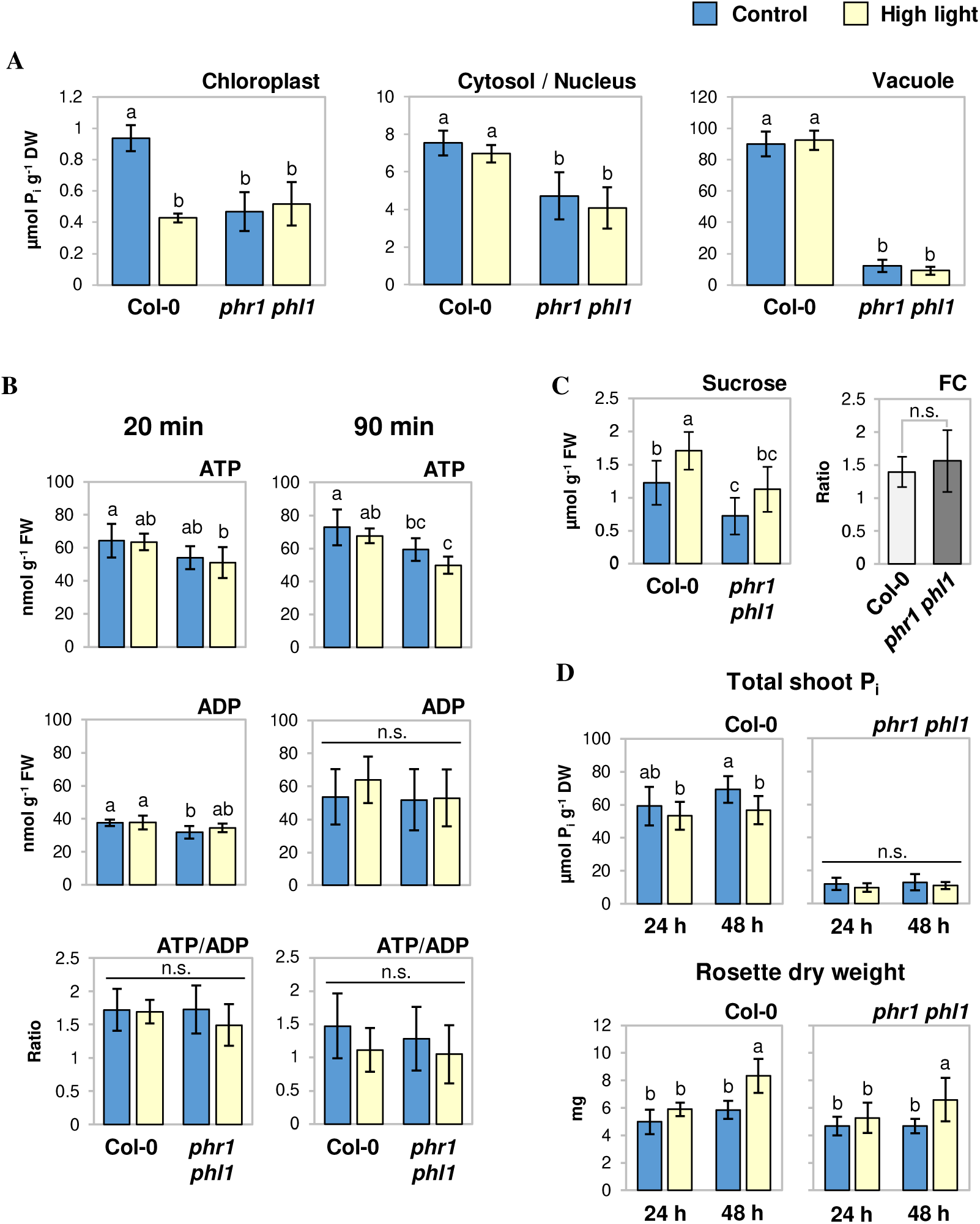
High light affects cellular P_i_ pools and sucrose levels. **A-C**, Plants were grown and treated as described in Fig. 2B. **A**, Subcellular P_i_ concentrations in WT and *phr1-1 phl1* mutant plants upon high light relative to dry weight (**DW**). Rosettes were harvested after 20 min of treatment. P_i_ contents were determined following non-aqueous fractionation of cellular compartments. *n* = 4 (*phr1-1 phl1*) or 5 (Col-0) independent experiments. **B,** Changes in ATP/ADP levels after shift to high light relative to fresh weight (**FW**). *n* = 6 plants from 3 (90 min) or 4 (20 min) independent experiments. **C**, Sucrose levels upon 20 min of high light; *n* = 8 plants from 4 independent experiments; absolute values and fold changes (**FC**, high light/control) are shown. Significant difference in FC was tested using Student‘s *t* test with 2-tailed distribution, unpaired with unequal variance. **D,** High light affects total shoot P_i_ and growth rates on the medium-term. WT and *phr1-1 phl1* mutants were grown for 4 weeks as described for Fig. 2B before light intensity was shifted to 450±30 µmol m^-2^ s^-1^ (high light) at 4 h after onset of the photoperiod. Control plants were kept under growth light conditions (70±5 µmol m^-2^ s^-1^). Material was harvested after 24 and 48 h. Total P_i_ contents in rosettes relative to DW and average rosette DWs are depicted; *n* = 10 plants from 5 individual experiments. All bars show means ± standard deviations. Except for FC shown in **C**, all statistical analysis were performed using 2-factor ANOVA with Tukey HSD post-hoc test; *P* < 0.05; n.s., not significant.

PHR1(-like) regulation by inositol pyrophosphates relates to the metabolism of these signaling compounds in the cytosol (Wild *et al*., 2016) which depends on both P_i_ and ATP levels (Zhu *et al*., 2019; Riemer *et al*., 2021). Given that cytosolic P_i_ concentrations were unchanged by 20 min high-light treatments, we asked if light increase affected cellular ATP levels. While *phr1-1 phl1* double mutants exhibited a tendency towards lover ATP levels that was more pronounced upon high-light treatment, light increase did not significantly affect ATP or ADP levels in the WT at 20 or 90 min of treatment (Figure 3B). Hence, activation of PHR1(-like) transcription by short-term high light appears to be independent of cytosolic P_i_ and ATP levels.

It has been reported that P_i_ starvation responses are modulated by sugar availability (Franco-Zorrilla *et al*., 2005; Müller *et al*., 2005, 2007; Karthikeyan *et al*., 2007). Since sugars are also involved in the signaling of photosynthetic status (Häusler *et al*., 2014; Li & Sheen, 2016; Zirngibl *et al*., 2023), we tested if short-term high-light treatment was sufficient to induce changes in sugar levels in leaves. At the timepoint tested, i.e. 80 min after onset of the photoperiod, contents of sucrose under control conditions were found to be lower in leaves of *phr1-1 phl1* as compared to the WT (Figure 3C). 20 min high-light treatment resulted in a further increase in sucrose (Figure 3C), but not glucose or fructose (Supporting Information Fig. S6) levels in the WT. A similar fold-change of sucrose contents was observed in *phr1-1 phl1* mutant leaves upon high light, although the increase in absolute levels was not significant (Figure 3C). Thus, sucrose levels respond to high-light treatment on a very short timescale, suggesting that they might act as early signaling compounds, and potentially might also affect light-dependent P_i_ starvation responses.

Analysis of transcriptomic data revealed upregulation of both P_i_ and nitrogen starvation responses upon medium-term (24 h) high light (Figure 2A). To address if this was attributable to enhanced nutrient requirements caused by high light, we determined total P_i_ contents of rosettes subjected to 24 and 48 h of light increase. In fact, shoot P_i_ levels of WT plants were slightly reduced within 48 h of high light as compared to control light conditions (Figure 3D). This correlated with significantly enhanced plant biomass measured upon 48 h of high light compared to control treatments (Figure 3D), suggesting that light increase stimulated growth which was not immediately satisfied by additional nutrient acquisition. In contrast to WT plants, P_i_ contents in leaves of *phr1-1 phl1* mutant plants remained at low levels independently of the light treatment (Figure 3D).

Together, our results indicate that induction of P_i_ starvation responses upon light increase occurs in at least two phases: Short-term (20-45 min) responses are triggered by a signal independent of cytosolic/nuclear P_i_ concentrations. This short-term signal communicates chloroplast P_i_ levels to the nucleus, and/or potentially involves cellular sucrose levels. In the medium term however, a shift to higher light intensities causes cellular P_i_ depletion through an imbalance of growth and nutrient uptake.

### Carbohydrate accumulation is affected in *phr1 phl1* mutants

We asked if the induction of P_i_ starvation gene expression was relevant for the acclimation of photosynthetic processes to increased radiation. To this end, we first performed pulse-amplitude modulated (PAM) chlorophyll fluorometry on rosette leaves subjected to increased light. The efficiency of photosystem II photochemistry (ΦPSII) dropped to the same extent in both WT and *phr1-1 phl1* mutants upon exposure to increased illumination for 125 min (Figure 4A). After 125 min of treatment, also the maximum efficiency of PSII (F_v_/F_m_) had decreased significantly in the stressed plants of both genotypes (Figure 4B). As for ΦPSII, no difference was observed for F_v_/F_m_ between WT and *phr1-1 phl1*, indicating that thylakoid reactions were not affected in *phr1-1 phl1* mutants.

**Figure 4.**
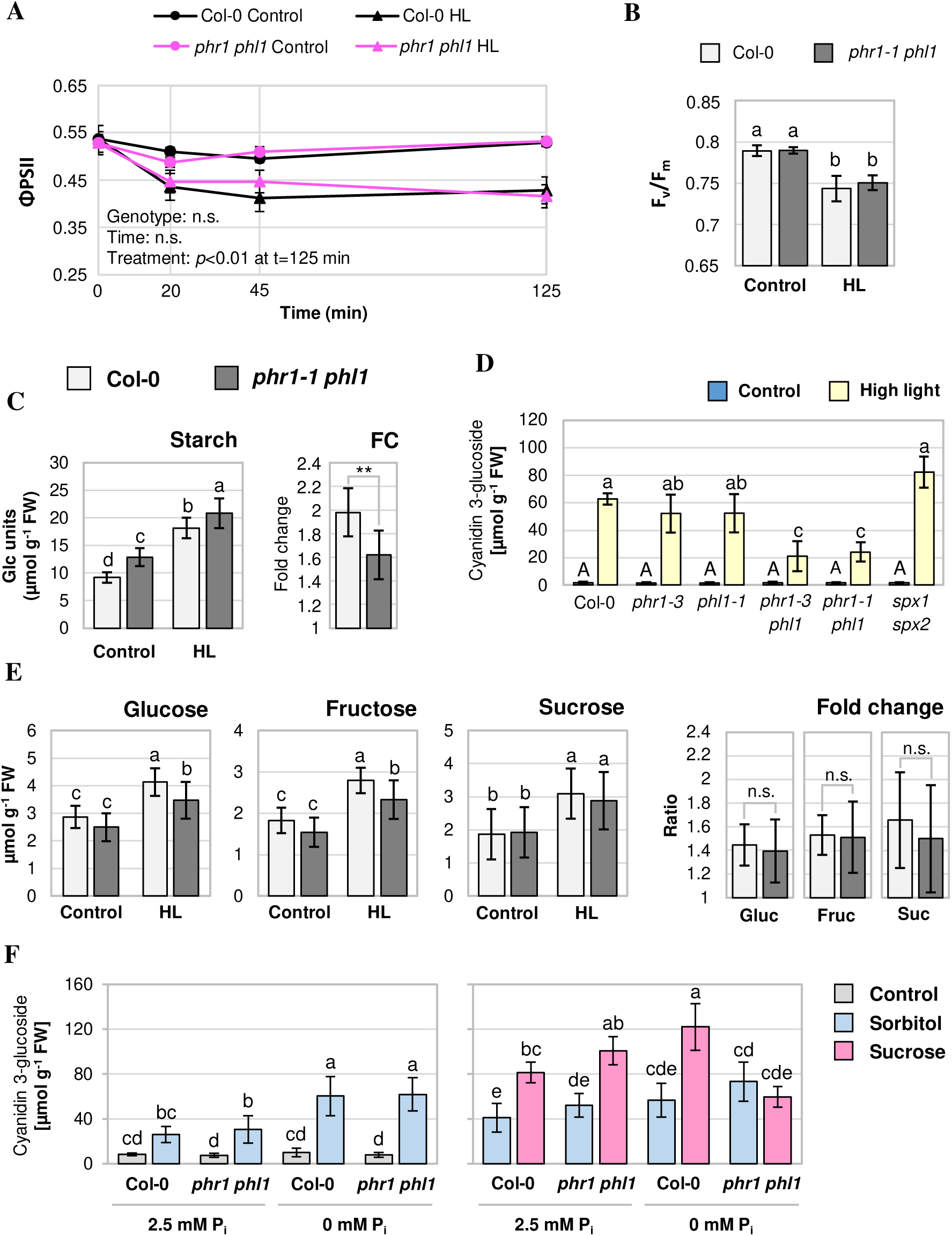
*phr1-1 phl1* mutants show altered carbohydrate responses. Unless specified, plants were grown and treated as described for Fig. 2B. Data points and bars depict means ± standard deviations. **A, B**, PAM measurements were performed to determine the effect of high light (**HL**) on chlorophyll a fluorescence parameters. *n* = 3 independent experiments (representing means of each 3 leaves measured). **A,** Quantum yield of photosystem II photochemistry (**ΦPSII**) in light-adapted leaves after 20, 45, and 125 min of treatment. Significance was tested using One-way ANOVA with Tukey HSD follow-up test and Bonferroni alpha correction for contrasts; n.s., not significant. **B,** Plants subjected to high-light stress were dark-adapted for 20 min after 125 min of stress duration to determine the maximum efficiency of photosystem II (F_v_/F_m_). 2-factor ANOVA with Tukey HSD post-hoc test; *P* < 0.05. **C, E**, Changes in carbohydrate levels in WT (Col-0) and *phr1-1 phl1* mutants after 90 min of high-light treatment. 2-factor ANOVA with Tukey HSD post-hoc test; *P* < 0.05. Fold changes (**FC**) of sugar/starch contents were calculated for each genotype as HL/mean of control. Statistical analyses of FC were performed with Student‘s *t* test with two-tailed distribution, unpaired with unequal variance. Significant differences are indicated as ** *P* < 0.01, n.s. not significant. **C**, Starch levels relative to fresh weight (**FW**); *n* = 9 plants from 3 independent experiments. **D**, Anthocyanin (cyanidin 3-glucoside) levels after 3 days of high-light exposure. Plants were grown for 32 days until onset of the treatment. Leaf material was harvested after additional 70 h. Pigment contents were calculated relative to FW. *n* = 3 independent experiments; statistical analyses were conducted for each treatment group using One-way ANOVA with Tukey HSD follow-up test and Bonferroni alpha correction for contrasts, *P* < 0.01. **E**, Levels of glucose, fructose, and sucrose after 90 min of high-light treatment. *n* = 11-12 plants from 4 independent experiments. **F**, *phr1-1 phl1* mutants are defective in sugar-induced anthocyanin production under P_i_-limiting conditions. Seedlings were grown on nutrient-rich media (no sucrose or sorbitol added) for 7 days under a 16-h light regime before transfer to media with either 0 or 2.5 mM KH_2_PO_4_ added and containing either no additives (control), 90 mM sucrose, or 160.7 mM sorbitol (same osmotic strength). Seedling shoots were harvested for anthocyanin determination after 67 h of growth on differing media. *n* = 6 pools of seedling shoots from 3 independent experiments; 2-factor ANOVA with Tukey post-hoc test; *P* < 0.05.

Next, we determined the changes in carbohydrate contents that occurred within 90 min upon an increase in photon flux density from 70±5 µmol m^-2^ s^-1^ to 450±30 µmol m^-2^ s^-1^ in rosette leaves of 5-weeks-old WT and *phr1-1 phl1* mutant plants. As shown in Figure 4C, starch contents under control conditions were higher in *phr1-1 phl1* mutants than in the WT. This may be related to the fact that *phr1-1 phl1* mutants manifest lower P_i_ levels in the chloroplasts (Figure 3A), a condition that allosterically activates AGPase (Figueroa *et al*., 2022). High-light treatment caused an increase in starch contents in both WT and mutant (Figure 4C). However, the fold change of starch content upon high light was lower in *phr1-1 phl1* than in WT plants. This was accompanied by slightly lower levels of monosaccharides (glucose and fructose), but not sucrose, in the double mutant upon 90 min of light increase (Figure 4E). Notably, differences in sucrose levels that were seen between WT and *phr1-1 phl1* under control growth at 80 min after the onset of the photoperiod (Figure 3C), were not detected at 150 min into the photoperiod (Figure 4E), indicating that sucrose levels are disturbed in *phr1-1 phl1* during the early morning.

In conclusion, carbon fixation and/or allocation is slightly compromised in *phr1-1 phl1* mutants upon high light, while WT-like sucrose levels are maintained during the later course of the day, independent of light conditions.

### PHR1/PHL1 are involved in high-light-mediated anthocyanin accumulation

Since both P_i_ starvation and high-light stress trigger anthocyanin production, we next analyzed the role of *PHR1/PHL1* signaling in anthocyanin accumulation during high-light acclimation. As shown in Figure 4D, anthocyanin contents of *phr1 phl1* leaves after 3 days of high-light exposure were significantly lower than in WT leaves or in the single mutants *phr1-3* and *phl1*, while loss of *SPX1/2* function did not have any statistically significant effect. PHR1 has been implicated in flavonoid biosynthetic gene induction under low P_i_ (He *et al*., 2021; Liu *et al*., 2022). We did not detect significant differences in the induction of *LDOX*, *DFR*, *PAP1*, or *MYB111* transcripts between WT and *phr1-1 phl1* mutants at 24 or 48 h of light increase (Supporting Information Fig. S7). Interestingly however, *PAP2* expression was markedly upregulated in *phr1-1 phl1* compared to the WT at 24 h of high light (Supporting Information Fig. S7), indicating that the regulation of anthocyanin accumulation is disturbed at a different level than biosynthetic gene expression in *phr1-1 phl1*.

It has been reported that high-light induced anthocyanin biosynthesis depends on sucrose signaling (Zirngibl *et al*., 2023). Given that sucrose levels were similar between WT and *phr1-1 phl1* at 90 min of high light (Figure 4E), we reasoned that sugar-mediated anthocyanin production might be impaired at the signaling level in *phr1-1 phl1*. To address this, we exposed 7-days old seedlings grown on nutrient-rich media to sucrose- or sorbitol-containing media with different P_i_ supplies for 67 hours. As depicted in Figure 4F, 3 days of mere P_i_ depletion were not sufficient to trigger measurable anthocyanin production in either genotype. When sorbitol was added to the media, WT and *phr1-1 phl1* seedlings accumulated anthocyanins to the same level (Figure 4F), indicating that WT and *phr1-1 phl1* are indistinguishable in terms of anthocyanin production upon osmotic stress. Crucially, when sorbitol containing media were compared to media containing sucrose at same-osmotic effect concentrations, *phr1-1 phl1* mutants failed to produce anthocyanins above the level of the osmotic effect when the growth medium lacked P_i_ (Figure 4F). Thus, *phr1-1 phl1* mutants are specifically defective in sucrose-mediated anthocyanin production already at early stages of P_i_ depletion.

Taken together, *PHR1/PHL1* functions are required for adequate high-light mediated anthocyanin accumulation, which might be related to a role in sucrose signaling upon cellular P_i_ restriction.

### *SRG3* expression is required for lipid homeostasis upon light increase

Among the *PHR1/PHL1*-dependent transcripts analyzed under short-term high light, *SRG3* showed particularly strong upregulation (Figure 2B), and it has been suggested that the *SRG3* promoter is directly activated by PHR1 (Cheng *et al*., 2011). To test this hypothesis, we performed promoter-transactivation assays in mesophyll protoplasts. PHR1 was able to induce the expression of a luciferase reporter fused to the promoters of *SRG3*, *MGD3*, *GPT2*, and *SPX1* (lacking the naturally occurring *Nco*I restriction site, *proSPX1^GC^*) (Figure 5A). Notably however, the responsiveness of *proGPT2* to *PHR1* expression was considerably lower than observed with the constructs harboring *proSPX1^GC^, proSRG3* or *proMGD3* upstream of the reporter CDS. This finding was consistent with the observation that *GPT2* was induced independently of *PHR1/PHL1* under high light (Figure 2B), and suggests that *SRG3*, like *MGD3* and *SPX1*, is a veritable target gene of PHR1.

**Figure 5.**
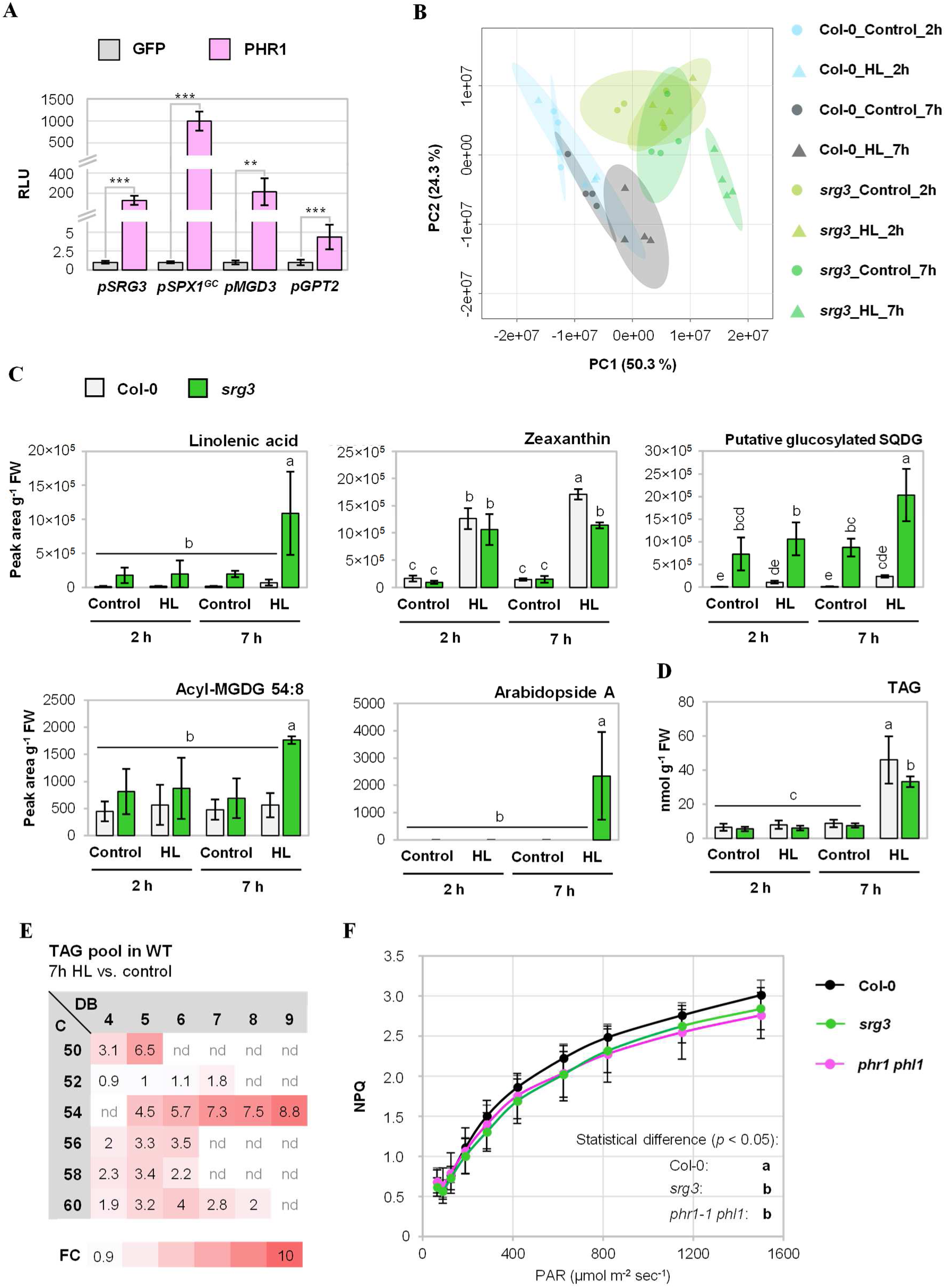
Effect of *srg3* mutation on the lipid composition upon light increase. **A,** Promoter transactivation assay in mesophyll protoplasts showing that the promoters of *SPX1*, *MGD3*, and *SRG3* are activated by PHR1. Promoter sequences were fused to the firefly *LUC* reporter gene (*proSRG3::LUC_Firefly_*; *proSPX1^GC^::LUC_Firefly_; proMGD3::LUC_Firefly_*, *proGPT2::LUC_Firefly_*) and co-transfected with either *PHR1* or *GFP*. Luminescence units were normalized to the luminescence of *Renilla LUC* which was additionally co-expressed from the Cauliflower mosaic virus *35S* promoter, and calculated relative to the luminescence level of GFP-expressing samples. **RLU**, relative luminescence level. Results for each promoter construct are from independently triplicated experiments. Means ± standard deviations; *n =* 6 from 3 independent experiments. One-way ANOVA with Tukey HSD follow-up test and Bonferroni alpha correction for contrasts; \*\*\**P* < 0.001, \*\**P* < 0.01. **B-E**, Plants of WT (Col-0) and *srg3* genotype were grown and treated as described for Fig. 2B, but leaves were harvested at 2 and 7 h of treatment. Lipid species were analyzed using LC-MS. **B**, Principal component analysis of samples from 2 h and 7 h high-light (**HL**) treated or control plants of WT and *srg3* genotype. **C**, Levels of *srg3*-dependent HL-responsive lipid features normalized to leaf fresh weight (**FW**). *n* = 4 independent experiments, except for acyl-MGDG: *n* = 3-4. **D, E,** *n* = 3-4. **D,** Contents of triacylglycerols (**TAG**) in WT and *srg3* leaves after high-light treatments. **E**, Changes in TAG pools upon 7 h of light increase (7 h HL vs. 7 h control) in the WT. Fold changes (Col-0_HL_7h/Col-0_C_7h) are indicated by color code and were calculated for TAG species sorted by the total number of carbon atoms of the acyl chains (**C**) and the total number of double bonds (**DB**). **nd**, not detected. **F**, Light response of the NPQ parameter measured by PAM chlorophyll fluorometry. Plants of WT (Col-0), *srg3*, and *phr1-1 phl1* genotype were grown for 40 days in an 8-h light regime at 70±5 µmol m^-2^ s^-1^ and dark-adapted for ≥30 min before measurement. NPQ was plotted against the fluence rate of photosynthetically active radiation (**PAR**). *n* = 3 independent experiments representing means of 2 plants. **C-F**, 2-factor ANOVA with Tukey HSD post-hoc test; *P* < 0.05.

We next asked whether PHR1-mediated induction of *SRG3* was relevant for lipid metabolism upon light increase, given the proposed role of SRG3 in phospholipid degradation pathways. To answer the question, we compared the lipid profile of a four-week-old WT and *srg3* loss-of-function mutant (Cheng et al., 2011) grown in soil, which had been exposed to high light for two or seven hours. The two-hour exposure was an early timepoint of acclimation, close to the onset of gene expression, while the seven-hour exposure corresponded to the end of the light period to allow for a stronger metabolic response. Total lipid extracts of shock-frozen leaf material were analyzed by ultra-performance liquid chromatography coupled to a quadrupole time-of-flight mass spectrometer equipped with electrospray ionization source. The first principal component of the PCA analysis (Figure 5B) on the aligned 2478 lipid features (Supporting Information Table S1) accounted for 50 % of the total variance, suggesting differences in the lipidome of *srg3* compared to the WT. Therefore, we profiled the main structural lipids monogalactosyl diacylglycerols (MGDG), digalactosyl diacylglycerols (DGDG), sulfoquinovosyl diacylglycerols (SQDG), phosphatidylcholines (PC), phosphatidylglycerols (PG) and phosphatidylinositols (PI) in WT and *srg3* after 2 h and 7 h of light stress. Their total levels were comparable (Supporting Information Fig. S8A), which is consistent with the results obtained for *srg3* mutants under P_i_ starvation conditions (Cheng *et al*., 2011). Notably, the levels of MGDG(34:1), DGDG(34:1) and DGDG(34:2) were slightly higher in both WT and *srg3* after 7 h of high light (Supporting Information Fig. S9), whereas the levels of the phospholipids determined were comparable (Supporting Information Fig. S10). In addition, profiling of sphingolipids revealed non-significant higher levels of ceramides in *srg3* compared to WT in all four investigated conditions, but hydroxy ceramides and glucosyl ceramides were WT-like in the mutant (Supporting Information Fig. S8B).

We also performed lipid profiling of acyl-MGDGs and Arabidopsides in the extracts, as galactolipids can be acylated or oxygenated (Ibrahim *et al*., 2011). We could only identify acyl-MGDG-54:8 and Arabidopside A (1-OPDA, 2-dnOPDA MGDG), and their levels were significantly higher in *srg3* after 7 h of high light stress compared to the other groups (Figure 5C). This result encouraged us to perform ANOVA on the annotated 2478 lipid features (Supporting Information Table S2) in order to identify more *srg3*-dependent high-light responsive lipids. We first selected lipid features that were significantly different between WT plants treated with 7 h of high light and untreated plants. ANOVA revealed 81 lipid features (fold change ≥ 2, *P* ≤ 0.05) whose intensities were different after exposure to high light for 7 h (Supporting Information Table S2). Of the 81 lipid features that respond to high light in the WT, 18 were significantly different in *srg3* compared to WT after 7 h of high light (Supporting Information Table S3). One of them was identified as linolenic acid, whose levels were higher in *srg3* compared to WT after 7 h but not 2 h of high light (Figure 5C), like Arabidopside A and MGDG-54:8. Interestingly, a lipid feature was constitutively higher in *srg3* and accumulated upon high-light treatment (Figure 5C), which we could tentatively assign to a glycosylated SQDG species (SQDG-34:1+C_5_H_8_O_4;_ Supporting Information Table S3). The identification of this unknown lipid species will need to be verified in the future using tandem mass spectrometry and nuclear magnetic resonance spectrophotometry after purification.

One of the *srg3*-dependent metabolite features showing highest light-responsiveness was identified as zeaxanthin (Figure 5C; Supporting Information Table S3), a photoprotective carotenoid and component of the xanthophyll cycle (Demmig-Adams & Adams, 1996; Havaux & Niyogi, 1999). In *srg3*, zeaxanthin accumulation following 7 h of high light was reduced by one third as compared to the WT, indicative of a reduced capacity for light energy dissipation in chloroplast membranes of *srg3*. To further investigate this, we performed PAM chlorophyll fluorescence measurements and determined the light response of the non-photochemical quenching (NPQ) parameter in WT, *srg3*, and *phr1-1 phl1* mutant plants. Indeed, the extend of NPQ was slightly but significantly reduced in both the *srg3* and the *phr1-1 phl1* mutants compared to the WT (Figure 5F). Taken together, *SRG3* expression is required for maintaining fatty acid homeostasis and the establishment of light protection upon high-light stress.

### *SRG3* contributes to triacylglycerol accumulation upon light increase

The observation that linolenic acid accumulated in *srg3* leaves upon high light (Figure 5C) raised the question as to whether containment of free fatty acids was impaired by *srg3* mutation. One strategy to reduce the levels of cytotoxic fatty acids in plant cells is their incorporation into triacylglycerols (TAG) under stress, including P_i_ starvation (Pant *et al*., 2015; Yang & Benning, 2018). We therefore quantified total TAG levels and found a pronounced accumulation after 7 h of light treatment in the wildtype (Figure 5D). In *srg3* mutants, high-light induced TAG accumulation was significantly lower. The differences between WT and mutant were due to reduced accumulation of TAG species with long-chain fatty acids exhibiting higher degree of desaturation (54:6, 54:7, 54:8, 54:9) in *srg3* (Supporting Information Fig. S12), which would be accordant with linolenic acid esterification. TAG species with polyunsaturated 54-carbon acyl chains also showed the highest changes in the WT upon high light (Figure 5E), a pattern that is reminiscent of heat-stress induced TAG accumulation (Mueller *et al*., 2015). To control if high light caused a heat response in the treated plants, we analyzed transcript levels of *HEAT SHOCK PROTEIN 70* (*HSP70*) and *HSP18.2*, previously used to discriminate light and heat responses (Huang *et al*., 2019). High light under our experimental conditions caused accumulation of *HSP70* and, to a lesser extent, *HSP18.2* transcripts which was mostly confined to the 45 min timepoint of treatment (Supporting Information Fig. S13). While *HSP70* induction upon high light was also seen by Huang *et al*. (2019), *HSP18.2* transcript accumulation is indicative of a mild heat response under the conditions used in this study. Thus, we cannot fully exclude that TAG accumulation observed upon high-light treatments was triggered by higher temperature at the leaf surface. Nevertheless, the late appearance of TAG enrichment after 7 hours of high-light treatment, as opposed to the moderate and transient induction of *HSP18.2* at 45 min of treatment, suggests that the observed changes might indeed be triggered by high-light treatment independently of temperature effects. Our analysis of TAG accumulation in *srg3* mutant plants furthermore suggest that *SRG3* gene function is required for normal occurrence of stress-induced TAG enrichment.

## Discussion

P_i_ is intimately linked to photosynthetic metabolism, both as a substrate for photophosphorylation, and as a potent kinetic regulator of carbon assimilation (Furbank & Lilley, 1980; Stitt *et al*., 1983; Preiss, 1984; Hendriks *et al*., 2003; Marcus *et al*., 2005). Additionally, P_i_ starvation responses affect the photosynthetic machinery of plants (Giersch & Robinson, 1987; Fredeen *et al*., 1990; Liu *et al*., 2024) which is partially achieved at the transcriptional level (Wu *et al*., 2003; Morcuende *et al*., 2007; Bustos *et al*., 2010; Nilsson *et al*., 2012). Nevertheless, it has remained elusive whether changes in the metabolic P_i_ demand of the chloroplast relay to the nuclear P_i_ starvation response. The observations presented herein establish that the P_i_ response machinery involving the transcription factors PHR1 and PHL1 is activated by an increase in photosynthetic activity, and we provide evidence that *PHR1/PHL1* contribute to the rapid metabolic adjustment during changes in light intensity (Figure 6).

**Figure 6.**
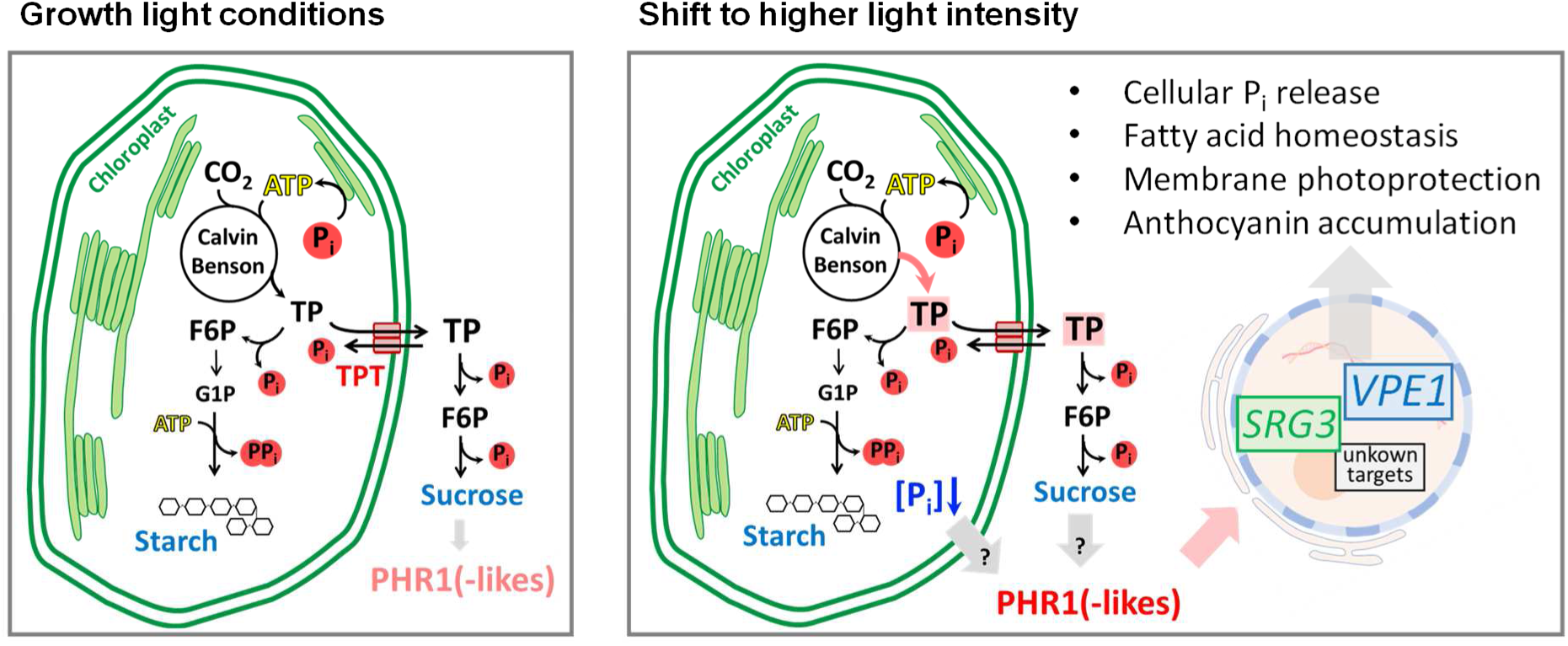
Model of P_i_ signaling under light increase. **Left**, Under normal growth light conditions, production of triose phosphates in the Calvin Benson cycle and their utilization in downstream processes such as starch and sucrose biosynthesis are balanced, and the rate of P_i_ recycling matches its consumption by the chloroplast metabolism (Stitt *et al*., 1987; McClain & Sharkey, 2019). **Right**, Upon shift to higher fluence rates, phosphorylated photosynthetic intermediates accumulate (Borghi *et al*., 2019), resulting in a decrease of chloroplast P_i_ concentrations. Chloroplast P_i_ levels or changes in the pools of photosynthetic products generate a signal that leads to PHR1(-like) activation. This causes P_i_-starvation responsive gene expression including the induction of *SRG3* and *VPE1*, leading to P_i_ release from intracellular pools, and changes in lipid metabolism enabling the containment of free fatty acids and xanthophyll cycle operation, as well as anthocyanin accumulation in the medium-term. **TP**, triose phosphate; **F6P**, Fructose 6-phosphate; **G1P**, Glucose 1-phosphate.

### Increase in light intensity creates a transient low-P_i_ signal

Transcriptional responses to changes in light intensity can be observed on the timescale of minutes (Vogel *et al*., 2014; Huang *et al*., 2019), but the signaling cascades behind are incompletely elucidated. We noted that the transcriptomic changes upon a shift to high light overlap with those observed during growth under P_i_-limited conditions (Figure 2A). Interestingly, a previous study addressing intraspecies variation of the transcriptional response to increased irradiance recognized the induction of P_i_-starvation responsive genes upon high light in two out of three tested accessions of Arabidopsis, and PHR1-target gene induction coincided with photosynthetic efficiency of the ecotypes (van Rooijen *et al*., 2018). Among the high-light and low-P_i_ induced transcripts, we quantified the expression of *SRG3*, *SPX1*, and *VPE1* during the early phase of high-light acclimation and determined that these transcripts transiently accumulate already at 20 min of exposure to increased photon flux density (Figure 2B). Crucially, gene induction was absent in *phr1-1 phl1* double mutants, qualifying this short-term high-light response as dependent on nuclear low-P_i_ signaling. Hence, plant cells are capable of exploiting the transcriptional potential of the low-P_i_ response module in order to react to environmental stimuli, even under nutrient-replenished conditions. Notably, high-light mediated induction was not seen to the same extend for all PHR1 targets examined, as expression of *MGD3* and *PS2* were only marginal (Figure 2B). This suggests that short-term high light affects only a subset of the transcriptional P_i_ starvation response, potentially reflecting differential chromatin accessibility of the genes (Barragán-Rosillo *et al*., 2021).

How are PHR1 and PHL1 activated by an increase in light intensity? It has been established that photosynthetic activity, through the production of phosphorylated assimilates, reduces the P_i_ pool available for photophosphorylation (Sharkey, 1985; Walker & Sivak, 1986; Robinson & Giersch, 1987; McClain & Sharkey, 2019), but the effect on other cellular compartments than the chloroplast has remained obscure. Our analysis of subcellular P_i_ contents revealed a reduction of chloroplast P_i_ by more than one half upon an increase in light intensity of 20 min duration (Figure 3A). Meanwhile, no decline in the cytosolic/nuclear or the vacuolar fractions was observed, proving high capacity of the cell to control subcellular P_i_ distribution. This finding however argues against InsP8-dependent PHR1(-like) activation under these conditions. Given the large difference in vacuolar P_i_ levels compared to the cytoplasm, we cannot fully rule out that a difference in the P_i_ levels between treated and control cytosolic preparations was masked by minor vacuolar contaminations. It is also possible that a drop in cytosolic P_i_ levels sufficient to trigger a peak in PHR1(-like) activity occurs transiently prior to the sampling timepoint and therefore precedes transcript accumulation seen at 20 min of treatment. However, our finding is consistent with the assumption that P_i_ exchange between chloroplast and cytosol (Santarius & Heber, 1965), and even more so between cytosol and vacuole (Woodrow *et al*., 1984) are rather slow processes. Additionally, the rapid increase in sucrose levels that can be observed upon high light (Figure 3C) indicates that the capacity for P_i_ liberation via sucrose biosynthesis might be sufficient to balance cytosolic P_i_ levels against depletion through triose phosphate import.

Intriguingly, our observations would be coherent with the possibility that P_i_ levels of the chloroplast are directly sensed to promote PHR1(-like) activation upon high light. Evidence for a role of chloroplast P_i_ levels in the acclimation of the photosynthetic machinery was recently provided by Raju et al. (2024) who investigated stromal P_i_ levels of mutants defective in P_i_ transporters of the chloroplast envelope using a FRET-based P_i_ sensor. Here, P_i_ levels in the stroma of *tpt*, *pht2;1*, and double mutants correlated with ΦPSII and NPQ parameters, while proton motif force and proton efflux efficiency of the ATP synthase, as determined by electrochromic shift measurements, were not affected after 2 min of a dark-light transition. The latter finding is corroborated by our result that ATP levels of WT plants are unaffected upon light increase, although ATP synthesis in the chloroplast would be expected to be kinetically constrained at the P_i_ concentrations measured in our system (Sharkey, 1985; McClain & Sharkey, 2019). Chloroplast P_i_ acting as an original retrograde signal would serve as an explanation for high constitutive PHR1(-like) activation in the *adg1 tpt-2* mutant (Figure 1E). Here, metabolically available P_i_ levels are likely very low given that the mutant is defective in major pathways of triose phosphate utilization. Hence, bulk release of P_i_ in *adg1 tpt-2* might be restricted to glycolysis and vacuolar phosphatase activity acting on phosphorylated organic compounds such as phosphoenolpyruvate (Ohnishi *et al*., 2018). This would cause unequal intracellular pool sizes of free P_i_, characterized by high vacuolar P_i_ stores (Figure 1D), and particularly low P_i_ levels in the chloroplast. It will be insightful to examine the subcellular distribution of P_i_ in the *adg1 tpt-2* mutant during future studies in order to elucidate the function of chloroplast P_i_ in retrograde signaling, and in the pleiotropic phenotypes of the *adg1 tpt-2* mutant.

### PHR1/PHL1 act in the network of sugar signaling

Blocking starch production in the *phr1-3 phl1* double mutant by introduction of the *adg1-1* allele substantially restricted growth (Figure 1B, C, F). In line with this, *phr1-1 phl1* mutants over-accumulated starch at the adult stage (Figure 4C), indicating that starch biosynthesis aids metabolism when cellular P_i_ levels are chronically depleted as it is observed in *phr1 phl1* mutants (Figure 3A, D). In fact, it is estimated that the bulk of P sequestered in photoassimilates is liberated via starch and sucrose biosynthesis (McClain & Sharkey, 2019). In line with the assumption that chloroplast P_i_ reserves are exploited through photosynthetic P_i_ consumption, growth restriction of *phr1-3 phl1*, *adg1-1*, and particularly *adg1 phr1 phl1* seedlings was more pronounced under high-light growth conditions (Figure 1F). However counter-intuitively, this was alleviated by an additional carbohydrate source supplied via the growth medium (Figure 1F), suggesting that metabolic alterations in these mutants extend beyond an insufficient capacity for P_i_ recycling for photoassimilation. It has been concluded from the studies on *adg1 tpt-2* mutants that sugars act as retrograde signaling molecules communicating photosynthetic status to the nuclear genome (Heinrichs *et al*., 2012; Häusler *et al*., 2014; Schmitz *et al*., 2014). Like in *adg1 tpt-2* (Schmitz *et al*., 2014), acclimation to high-light growth conditions might be impaired in *adg1 phr1 phl1*, and to a lesser extend in *phr1 phl1* mutants, resulting from a depletion of carbohydrates that occurs in the long-term of light exposure. In fact, already under the moderate growth light conditions, we noted time-of-day dependent deficits in sucrose levels in adult *phr1-1 phl1* plants (Figure 3C, 4E), similar to *adg1 tpt-2* mutants that lack diurnal peaks in sucrose concentrations (Schmitz *et al*., 2012). Moreover, *phr1 phl1* mutants exhibit a phenotype of early leaf senescence (Bustos *et al*., 2010; Wang *et al*., 2023) that is reminiscent of the symptoms caused by carbohydrate depletion. Interestingly, Schmitz et al. (2014) also noted that *adg1-1* mutation was associated with transcriptomic changes affecting lipid metabolism as we report here for the *PHR1/PHL1*-mediated high-light response. Finally, *phr1 phl1* mutants were specifically insensitive towards sucrose in terms of anthocyanin production under early P_i_-depleted growth conditions (Figure 4F). We therefore propose a function of PHR1/PHL1 in sugar signaling, thus contributing to the adjustment of the C/P balance.

### PHR1/PHL1 promote acclimation to higher light intensities

Our analysis of the *PHR1/PHL1* dependent transcriptional response to high light focused on *SRG3*, *SPX1*, and *VPE1*. *VPE1* induction, putatively encoding a vacuolar efflux transporter (Xu *et al*., 2019), immediately suggests itself as a strategy for intracellular P_i_ redistribution in favor of the cytosol by exploitation of the vacuolar P_i_ stores. This would serve to provide additional P_i_ to metabolic pathways, which seems adequate since light increase represents a condition that accelerates growth rate and implies higher nutrient demand in the mid-term (Figure 3D). In contrast, *SPX1* as a highly responsive P_i_ starvation marker gene (Wang *et al*., 2023) functions in negative feed-back of P_i_ starvation responses (Puga *et al*., 2014). Such fine-tuning of PHR1(-like) activation under light increase seems appropriate given the fluctuating nature of this environmental factor.

The pronounced PHR1/PHL1 dependent induction of *SRG3* was further investigated within this study. The mechanistic contribution of SRG3 to P_i_ homeostasis is indistinct, and it has been reported in other plant species that GDPD isoforms play a role in phospholipid homeostasis (Dang *et al*., 2025), but also in hormone signaling and the modulation of root morphology upon P_i_ starvation (Mai *et al*., 2021; Hu *et al*., 2024; Verma *et al*., 2025). We demonstrated here that the *SRG3* promotor is activated by over-expression of *PHR1* in mesophyll protoplasts (Figure 5A), indicating that it constitutes a true target of the transcription factor. To elucidate a potential physiological function of *SRG3* expression for lipid metabolism in the light acclimation response of shoots, we set out to identify lipid features that were altered by *srg3* mutation upon high-light treatment using an untargeted approach. Comparing WT and *srg3* rosette leaves, we detected differences in the lipid fingerprints particularly after longer treatment duration (Figure 5B). Among the molecules that over-accumulated in *srg3* mutants in a high-light dependent manner, we identified linolenic acid, the galactolipid derivatives acyl-MGDG (54:8), and Arabidopside A, as well as a molecule that was constitutively upregulated in *srg3* and putatively represents glucosylated SQDG (Figure 5C). While the latter lipid species has to our knowledge not been reported before, accumulation of the remaining species draws the picture that *srg3* mutants experience stress during acclimation to higher light intensities. Both acyl-MGDG and Arabidopside accumulation have been reported to be associated with stress conditions (Buseman *et al*., 2006; Vu *et al*., 2014; Nilsson *et al*., 2016), and it was suggested that the pool of acyl-MGDG increases upon stress to sequester oxidized fatty acids (Vu *et al*., 2014). Moreover, Arabidopside A has been ascribed a role in the regulation of chlorophyll degradation during senescence processes (Weber, 2002; Hisamatsu *et al*., 2006). Thus, the lipid fingerprint of *srg3* upon light increase implies that the gene might be required to maintain the integrity of photosynthetic membrane components upon changes in light availability. The accumulation of linolenic acid in leaves of light-stressed *srg3* plants furthermore indicates that the processing of fatty acids is disturbed in the mutant. After 7 h of high light, we observed an increase in galactolipid species with low degree of desaturation in WT and mutant plants (Supporting Information Fig. S9), an aspect of short-term high-light acclimation that has been reported before (Burgos *et al*., 2011). Such membrane remodeling involves lipid degradation and can be assumed to generate free fatty acids with higher degree of desaturation, including linolenic acid. Hence, SRG3 might fulfill a function in the containment of such products derived from membrane remodeling. The product of SRG3 activity, glycerol-3-phosphate (Cheng *et al*., 2011), is a substrate for membrane lipid biosynthesis, as well as for TAG production (Bates, 2016), both of which also require fatty acid precursors. To test whether any of these pathways is constrained in *srg3*, giving rise to higher linolenic acid levels, we quantified the pools of predominant membrane lipids and TAGs. While the levels of major lipid classes were unaffected by *srg3* mutation (Supporting Information Fig. S8), we detected TAG accumulation in the WT after 7 h of light increase which was slightly reduced in *srg3* mutants (Figure 5C). Consistent with a potential involvement of SRG3 in plastoglobule formation, TAG profiles revealed that the species with strongest accumulation under our high-light conditions also contained linolenic acid (Figure 5E, Supporting Information Fig. S12). Notably, linolenic acid is also a component of the acyl-MGDG (54:8) species which was enriched in *srg3* upon high light, confirming high availability of this fatty acid in *srg3* under stress. Elevated levels of free fatty acids can be cytotoxic probably due to their amphipathic and detergent-like properties leading to the inhibition of photosynthesis and damage to PSII (Kunz *et al*., 2009). Thus, our results suggest that SRG3 contributes to the detoxification of free linolenic acid that is released during high light-induced membrane remodeling by incorporation of excess linolenic acid into TAGs. Notably, linolenic acid accumulation was previously also observed in Arabidopsis plants subjected to salinity stress, and this was connected to the control of ion homeostasis via the regulation of H^+^-ATPase activity (Han *et al*., 2017; Abdelrahman *et al*., 2021). Hence, instead of a direct involvement of SRG3 in the processing of fatty acids, linolenic acid accumulation in *srg3* might also be a secondary effect related to perturbed membrane remodeling.

Interestingly, a compound that accumulated upon light increase in the WT, but was reduced by *srg3* mutation, was identified as zeaxanthin (Figure 5C), a molecule that is involved in the protection against excess excitation energy as part of the xanthophyll cycle (Demmig-Adams & Adams, 1996; Havaux & Niyogi, 1999; Sacharz *et al*., 2017). In line with a function of both *SRG3* and *PHR1*/*PHL1* for proper operation of the xanthophyll cycle, assessment of the NPQ component of chlorophyll a fluorescence revealed deficits in both mutant genotypes (Figure 5F). It is well-known that the membrane lipid composition is important for the xanthophyll cycle, and both lipid class and fatty acid composition determine faithful operation of all components (Goss & Latowski, 2020). Hence, it is tempting to hypothesize that *SRG3* expression, induced by the nuclear low-P_i_ response machinery, might be required for the adjustment of the membrane environment to ensure proper function of the enzymes involved in the xanthophyll cycle. Strikingly, this might be directly related to the changes in chloroplast P_i_ levels seen under high light (Figure 3A), given the recent observations on NPQ levels of P_i_-transport mutants made by Raju et al. (2024) which were mentioned above. We therefore propose a signaling pathway originating from photosynthetic activity that engages PHR1 and PHL1 to induce *SRG3* expression for the acclimation of the photosynthetic membranes towards high light.

### Conclusions

High-light stress requires immediate responses to avert damage to cellular components. As a new mode of signaling excess photosynthetic activity, we describe a function of the low-P_i_ response machinery involving PHR1 and PHL1 transcription factors under nutrient replete growth conditions. Our results indicate that chloroplast P_i_ sequestration into phosphorylated photoassimilates, and/or the accumulation of these photoassimilates, create a retrograde signal resulting in the activation of PHR1 and PHL1 in the cytosol and nucleus. This serves to mobilize cellular P_i_, and to support acclimation responses including anthocyanin production, and the adjustment of lipid metabolism to ensure photoprotection and fatty acid homeostasis.

## Materials and Methods

### Plant material

All Arabidopsis plants used in this study are in the background of the Columbia-0 accession. The *phr1-3* (AT4G28610; SALK_067629C) (Nilsson *et al*., 2007) and *phl1* (AT5G29000; SAIL_731_B09) (Klecker *et al*., 2014) lines were described previously and the derived homozygous double mutant *phr1-3 phl1* was a kind gift of Angelika Mustroph (University of Bayreuth). The double mutant *phr1-1 phl1* (Bustos *et al*., 2010) was backcrossed to *phl1* in order to remove a transgene containing the *NPTII* cassette which was previously introduced for isolation of the *phr1-1* allele (Rubio *et al*., 2001). All experiments were performed with the homozygous double mutant *phr1-1 phl1* lacking the *NPTII* cassette. The double mutant *spx1 spx2* was described earlier (Puga *et al*., 2014). Knockout lines *adg1-1* (At5g48300) (Lin *et al*., 1988), *tpt-2* (At5g46110; SALK_073707.54.25.x) and *adg1 tpt-2* (Schmitz *et al*., 2012) were kindly provided by Rainer E. Häusler (University of Cologne). *gpt2-1* (AT1G61800; GK-454H06) and *srg3*/*gdpd1-1* (AT3G02040; SALK_087106) were obtained from The Nottingham Arabidopsis Stock Centre (NASC) and were described previously (Niewiadomski *et al*., 2005; Cheng *et al*., 2011). Primer sequences used for genotyping are listed in Table S4. Homozygous insertion mutants were verified using the following primer combinations: *PHR1* WT: PHR1_F/PHR1_R; *phr1-3* T-DNA: LBb1/PHR1_R; *PHL1* WT: PHL1_F/PHL1_R; *phl1* T-DNA: PHL1_F/LB3; *TPT* WT: TPT_F/TPT_R; *tpt-2* T-DNA: *tpt-2*_F/LBb1; *GPT2* WT: GPT2_F/GPT2_R; *gpt2-1* T-DNA: GK-LB/GPT2_R; *SRG3* WT: qSRG3_F/SRG3_R; *srg3* T-DNA: qSRG3_F/LBb1. To generate the *adg1 phr1 phl1* triple mutant, *adg1-1* was crossed to *phr1-3 phl1*. Homozygous *adg1-1* mutants were identified by screening for starch-free phenotypes by iodine staining at the end of the photoperiod. Homozygosity of the *phr1-1* allele was verified by Sanger sequencing (Macrogen Europe, Amsterdam) of the PCR product of PHR1_F/PHR1_R using primer PHR1_R.

### Cultivation conditions

Seeds were stratified for 64 hours at 4°C in the dark. For vegetative growth, plants were germinated and grown in an 8 h/16 h (light/dark) cycle and 23°C/22°C day/night temperature on soil consisting of seeding compost, universal soil, and vermiculite mixed in a 3:3:1 ratio. For high-light experiments, the plants were additionally fertilized once directly after pricking at the age of 10 days using a complete mineral mixture (Wuxal Super, Aglukon Spezialduenger, Duesseldorf, Germany) according to the manufacturer’s instructions. For seedling experiments, seeds were surface sterilized by chlorine gas exposure and transferred to solidified Arabidopsis growth media as described in the figure legends. Seedlings were grown under 16 h/8 h light/dark cycles. If not indicated otherwise, growth light was set to 90±10 µmol m^-2^ sec^-1^ (for seedling growth and for experiments shown in Figure 1B-E), or 70±5 µmol m^-2^ s^-1^ (before high-light treatments at 450±30 µmol m^-2^ s^-1^ of rosette plants), and monitored using an illuminance meter. Growth and treatments of rosette plants for high-light experiments were conducted in a Percival SE-41L plant growth incubator equipped with dimmable fluorescent lamps under temperature control.

### P_i_-limitation experiments

For P_i_-limitation experiments, seeds were germinated on medium containing MS salts at full strength together with 0.8 % (w/v) phytoagar (Duchefa Biochemie, Haarlem, The Netherlands) and 0.5 % (w/v) sucrose. After 5-10 days of growth (see figure legends), seedlings were transferred to P_i_ media (0.5 % (w/v) sucrose; 20 mM 2-(N-Morpholino)-ethane sulphonic acid; 2.5 mM KNO_3_; 1 mM MgSO_4_; 1 mM Ca(NO_3_)_2_; 2.5 mM KH_2_PO_4_; 25 µM Fe-EDTA; 35 µM H_3_BO_3_; 7 µM MnCl_2_; 0.25 µM CuSO_4_; 0.5 µM ZNSO_4_; 0.1 µM NaMoO_4_; 5 µM NaCl; 0.005 µM CoCl_2_; pH 6) solidified by 0.8 % (w/v) agar (adopted from Härtel et al. (2000)). For P_i_ depleting conditions, KH_2_PO_4_ was omitted, and Fe-EDTA was reduced to 10 µM in order to minimize low-P_i_ induced iron toxicity (Ward *et al*., 2008). Seedling shoots were harvested after 7-8 days of growth on P_i_ differing media (see figure legends). For the experiment shown in Figures 1F and Supporting Information Fig. S3, P_i_ medium was used containing either 0.25 mM KH_2_PO_4_/10 µM Fe-EDTA, 2.5 mM KH_2_PO_4_/25 µM Fe-EDTA, or 2.5 mM KH_2_PO_4_/25 µM Fe-EDTA +50 mM sucrose. For the experiment shown in Figure 4F, seedlings were germinated on MS salts at ½ strength without added sucrose and grown for 7 days before transfer to P_i_-differing conditions.

### Vector construction

Standard molecular cloning procedures were applied. Primer sequences used for cloning are listed in table S4. The firefly *LUCIFERASE* reporter plasmid *pBT10-proMGD3::LUC_Firefly_* and the *pHBTL-p35S::3HA-GFP* effector plasmid (Klecker *et al*., 2014), as well as the *pBT10-p35S::LUC_Renilla_* normalization plasmid (Bäumler *et al*., 2019) have been described elsewhere. For details on cloning *proSPX1^GC^, proSRG3*, *proMGD3*, *proGPT2,* and the *PHR1* effector construct, refer to the supplemental methods.

### Anthocyanin determination

Anthocyanin contents were determined with few changes as described in (Vandenbussche *et al*., 2007). In brief, 40-70 mg of seedling shoots or rosette leaf material were frozen in liquid nitrogen and ground in a bead mill (MM400, Retsch, Haan, Germany). The frozen powder was resuspended in 300 µl of methanol containing 1 % HCl. After vortex-mixing for 15 sec, cell debris was pelleted by centrifugation for 15 min at >14,000 g, 4°C. The supernatant was mixed with 200 µl water and 500 µl chloroform by vortexing for 15 sec. After 15 min centrifugation at 17,900 g, 4°C, the upper phase was diluted in methanol to measure the absorbance at 535 nm and 630 nm using a spectrophotometer (Specord 200 Plus, Analytik Jena, Jena, Germany).

### Measurements of starch, soluble sugars, and adenylates

Carbohydrates were determined by the Warburg optical test and adenylates were assessed based on luciferase activity. Both procedures were described previously (Mustroph *et al*., 2006), for details see supplemental methods.

### Determination of P_i_ contents

Free P_i_ was determined in leaf tissue based on the method described by (Ames, 1966) with some modifications partially based on (Sakuraba *et al*., 2018). For the experiment shown in Figure 3D, leaf material was lyophilized prior to extraction, and 50 µl extraction buffer per mg plant material was used. For details see supplemental methods.

### Non-aqueous fractionation of leaf material

For the analysis of subcellular P_i_ distribution, 12 plant rosettes (total of >1 g) were harvested into vials precooled by liquid nitrogen and dry ice, with an averaged harvesting time of 3 seconds (max. 7 seconds) per rosette. The plant material was ground to a fine powder and lyophilized. The freeze-dried powder was suspended in a mixture of tetrachlorethylene and n-heptane and fractionated across a non-aqueous density gradient as described before (Hernandez *et al*., 2023). Following fractionation, marker enzyme activities for plastids (alkaline pyrophosphatase), cytosol (UDP-glucose pyrophosphorylase), and vacuole (acidic phosphatase) were determined photometrically. Phosphate amounts were determined as described above for each fraction and correlated with the marker enzyme activities using the using the NAFalyzer app (https://github.com/cellbiomaths/NAFalyzer).

### RNA extraction and cDNA synthesis

Plant material was harvested under the respective experimental conditions and instantly frozen in liquid nitrogen. The tissue was ground in a bead mill, and RNA was extracted using Bioline TRIsure (Meridian Bioscience, Cincinnati, United States) according to the manufacturer’s instructions. The quality of the RNA was assessed by screening A_260_ and A_280_. For qRT-PCR analysis, 1 µg of RNA was treated with DNAse I (Thermo Fisher Scientific, Waltham, United States) according to product instructions before application in reverse transcription using oligo(dT)_15_ and RevertAid Reverse Transcriptase (Thermo Fisher Scientific). Complementary DNA was diluted by a factor of 50 for use as template in qPCR analysis. qPCR was performed in technical triplicates using SsoAdvanced Universal SYBR Green Supermix (Bio-Rad, Hercules, United States) on a CFX Connect Real-time System (Bio-Rad). Transcript levels were normalized to the levels of *PP2A* transcript and calculated as 1000·2^-ΔCT^. Primer sequences are listed in table S4.

### Protoplast isolation, transfection and transactivation assay

Leaf mesophyll protoplasts were isolated using the “Tape-Arabidopsis Sandwich” method (Wu *et al*., 2009) with minor modifications. For details, see supplemental methods.

### Chlorophyll fluorescence measurements

For the determination of the quantum yield of photosystem II photochemistry (*F*_m_′-*F*′)/*F*_m_′ (ΦPSII), as well as the maximum efficiency of photosystem II (*F*_v_′/*F*_m_′), and the NPQ parameter (or SV_N_; calculated by the Junior-PAM software using the Stern-Volmer equation for fluorescence quenching, Gilmore and Yamamoto (1991)), pulse-amplitude-modulated fluorescence measurements (PAM) of chlorophyll fluorescence were conducted using a Junior-PAM (Walz, Effeltrich, Germany). Here, to calculate chlorophyll fluorescence parameters using the saturating pulse method (Schreiber *et al*., 1986), one large rosette leaf was held between two magnets and subjected to 10 seconds of actinic light irradiance set to control light intensity (70±5 µmol m^-2^ sec^-1^), followed by a saturation light pulse.

### Lipid analysis

Lipid analysis was performed according to (Mueller *et al*., 2015). Progenesis QI (version 2.1, Waters) was used for data pre-processing and PCA was performed in the MetaboAnalyst platform (version 6.0, https://www.metaboanalyst.ca).

### Statistical analysis

Statistical analyses were performed with Microsoft Office Excel using the Real Statistics Resource Pack software (https://www.real-statistics.com). Two-factor ANOVA and hypergeometric testing were performed using RStudio (version 2024.09.1+394) available at https://www.r-project.org/.

## Supporting information

Supporting Methods and References

## Acknowledgements

We thank Stephan Clemens and Angelika Mustroph for helpful discussions and comments on the manuscript, as well as for providing infrastructure and equipment. We thank Susanne Berger for help with choosing experimental conditions used for lipid analyses.

## Author contributions

M.K., conceptualization; A.F., T.N., and M.K., methodology; L.A., A.F., T.N., and M.K., formal analysis; L.A., A.F., T.N., A.H., M.M., and M.K., investigation; M.K., writing–original draft; all authors, writing–review and editing; A.F. and M.K., visualization; M.J.M and M.K., supervision; M.K., project administration.

## Data availability

The source data of lipid analyses are available in the Supporting information of this article.

## Supplemental Data

**Supporting Information Fig. S1** Phenotypes of *adg1-1 phr1-3 phl1* (TM).

**Supporting Information Fig. S2** Expression of selected P_i_-starvation responsive genes depends on *PHR1/PHL1* under P_i_ depletion.

**Supporting Information Fig. S3** Root growth of *adg1 phr1 phl1* and *adg1 tpt-2* responds to exogenously applied sucrose.

**Supporting Information Fig. S4** *PHR1* transcript levels are not affected by short-time high-light exposure.

**Supporting Information Fig. S5** Changes in sugar and starch contents under P_i_ depletion.

**Supporting Information Fig. S6** Monosaccharide levels after 20 min of high light.

**Supporting Information Fig. S7** Anthocyanin biosynthetic and regulatory gene expression upon high light in WT and *phr1-1 phl1*.

**Supporting Information Fig. S8** Total levels of lipid classes in WT (Col-0) and *srg3* mutants upon shift to high light (HL).

**Supporting Information Fig. S9** Contents of glycosylglycerol lipid species in WT (Col-0) and *srg3* mutants upon shift to high light (HL) relative to fresh weight (FW).

**Supporting Information Fig. S10** Contents of glycerophospholipid species in WT (Col-0) and *srg3* mutants upon shift to high light (HL) relative to fresh weight (FW).

**Supporting Information Fig. S11** Levels of zeaxanthin and putative glycosylated SQDG determined in opposite ESI ion mode compared to Figure 5C.

**Supporting Information Fig. S12** Levels of triacylglycerol (TAG) species in WT (Col-0) and *srg3* mutants upon shift to high light (HL).

**Supporting Information Fig. S13** Levels of high-temperature responsive transcripts *HSP70* and *HSP18.2* in WT (Col-0) and *phr1-1 phl1* mutant plants upon high light treatment.

**Supporting Information Table S1** Annotated lipid features in WT (Col-0) and *srg3* mutants following a shift to high light.

**Supporting Information Table S2** Lipid features whose levels were significantly altered upon shifting to the WT (Col-0).

**Supporting Information Table S3** Heat-responsive lipid features whose levels differed significantly between WT (Col-0) and *srg3* mutants.

**Supporting Information Table S4** List of primer sequences used in this study.

**Supplemental Methods and References**

