## Supporting Methods and References for "PHR1 and PHL1 mediate rapid high-light responses and acclimation to high photosynthetic activity"

### Supplemental Methods

#### Vector construction

To generate the reporter plasmid *proGPT2::LUC$_{Firefly}$*, the *Bam*HI/*Nco*I fragment of *pBT10-proMGD3::LUC$_{Firefly}$* was ligated with the 5' upstream genomic sequence of *GPT2* (-1,501 bp from ATG). For *pBT10-proSRG3::LUC$_{Firefly}$* construction, the *SRG3* promoter region (-908 bp from ATG) was inserted into *pBT10-proMGD3::LUC* using *Bam*HI/*Nco*I restriction sites. *pBT10-proSPX1$^{GC}$::LUC$_{Firefly}$* was generated by replacing *proMGD3* in *pBT10-proMGD3::LUC$_{Firefly}$* by the 5' upstream genomic sequence of *SPX1* (-1,451 bp from ATG) using *Eco*RI/*Nco*I restriction sites. Given the occurrence of one *Nco*I site within the promoter sequence of *SPX1*, a G to C point mutation was introduced at -871 bp from ATG using primers proSPX1_mut_F and proSPX1_mut_R. Of note, the site of the G to C substitution does not affect any of the P1BS (GNATATNC) or P1BS-like (allowing two mismatches) motifs present in the *SPX1* promoter sequence. All promoter fragments were amplified from Col-0 gDNA.

To create the *PHR1* effector construct, the triple HA tag was amplified from a plasmid containing the *K2* cassette (Faden et al., 2016) using primers attB1-HA_F and PHR1-HA_R, and the *PHR1* sequence including introns and 3' UTR was amplified from Col-0 gDNA using primers HA-PHR1_F and attB2-PHR1_R, adding a linker sequence (encoding GSAG) to the *PHR1* 5' terminus. The fragments were subsequently stitched by PCR using primers attB1_F and attB2_R, and the PCR product was recombined into *pDONR201* (Invitrogen, Thermo Fisher Scientific, Waltham, United States) by a BP reaction using BP clonase II enzyme mix (#11789020, Thermo Fisher Scientific). *pDONR201-3xHA-PHR1* was used in LR reaction (LR clonase II; # 11791100, Thermo Fisher Scientific) with *pHBTL-p35S::3xHA-GW* (Ehlert et al., 2006), giving rise to *pHBTL-p35S::6xHA-$^{GSAG}$PHR1*.

##### Determination of starch, soluble sugars, and adenylates

For carbohydrate determination, 51-100 mg of rosette leaf material or seedling shoots were harvested under the respective growth conditions and instantly frozen in liquid nitrogen. Frozen plant samples were ground by a bead mill. 333 µl of 0.83 M perchloric acid was added to each sample, mixed well, and thawed on ice. Samples were centrifuged for 30 min at 15,000 *g*, 4°C. For the extraction of soluble sugars, the supernatants were mixed with 83.3 µl of 1 M bicine, and the mixture was immediately neutralized by addition of 50 µl of 4 M KOH. The extracts were pelleted for 10 min at 15,000 *g*, 4°C, and supernatants were stored at -80°C. For starch extraction, pellets from perchloric acid extraction were washed twice with 600 µl of 80 % ethanol before addition of 133 µl 0.2 M KOH and homogenization. The homogenates were incubated at 95°C for 1 h, followed by centrifugation at 15,000 *g* for 10-30 min. Supernatants were neutralized with 26.7 µl 1 M acetic acid. An extract volume of 50 µl was mixed with 100 µl of a 0.59 U/ml solution of amyloglucosidase (from *Aspergillus niger*, #A1602, Sigma-Aldrich, St. Louis, United States) in 50 mM sodium acetate, pH 5.0, and incubated overnight at 55°C.

To determine glucose contents, 20 µl of sample was mixed with 780 µl assay buffer (0.1 M imidazole; 5 mM $MgCl_2$; 2 mM nicotine amide adenine dinucleotide (#AE11.2, Carl Roth, Karlsruhe, Germany); 1 mM ATP disodium salt; pH 6.9) containing 2.8 U glucose-6-phosphate dehydrogenase (from *Leuconostoc*, #11079127, Roche, Basel, Switzerland). After addition of 0.5 U hexokinase (#11819031, Roche, Basel, Switzerland), absorbance at 340 nm was recorded continuously using a spectrophotometer (SPECORD 200 PLUS, Analytik Jena, Jena, Germany) until a plateau was reached. Likewise, for the subsequent measurements of fructose and sucrose contents, 0.2 U of phosphoglucose isomerase (from baker's yeast, #P5381-1KU, Sigma-Aldrich, St. Louis, United States) and 60 U of invertase (from baker's yeast, #I4504, Sigma-Aldrich, St. Louis, United States) were added sequentially. Differences in absorbance were used to calculate NAD consumption coupled to monosaccharide concentration.

##### Adenylate measurements

For the determination of adenylate levels, luciferine solution was prepared on the day prior to the measurement by mixing 50 ml of 25 mM HEPES·KOH, pH 7.75, with 120 mg bovine serum albumin, 171.6 g magnesium acetate, 31 µl of 65 mM DTT, and 1.323 ml of an aqueous solution of 7,56 mg/ml D-luciferine sodium salt (#102131, PJK, Kleinblittersdorf, Germany). This solution was mixed with 25 µl of 1 mg/ml luciferase (#102311, PJK, Kleinblittersdorf, Germany) in 0.5 M Tris acetate buffer (pH 7.5), and stored in the dark overnight. For ATP determinations, 20 µl of sugar extracts diluted in $H_2O$ by a factor of 10 were mixed with 200 µl of 25 mM HEPES·KOH, pH 7.75. For ADP measurements, ADP assay buffer was prepared on the day of measurement, by mixing 20 ml of 6.25 mM HEPES·KOH (pH 7.75) with 50 µl of 1 M magnesium acetate, 2 mg phosphoenolpyruvate, and 14 mg activated charcoal powder. After stirring for 5 min, the solution was paper filtrated to remove charcoal. The pellet obtained from 140 µl pyruvate kinase suspension (#P1506-1KU, Sigma-Aldrich, St. Louis, United States) was washed in 500 µl 3.2 M $(NH_4)_2SO_4$, and resuspended in the ADP assay buffer. 200 µl of this suspension was mixed with 20 µl of 10-fold diluted extracts of soluble sugars described above. For ATP and ADP measurements, the luciferase reaction was started by automated addition of 100 µl luciferine solution per 220 µl of extract-containing solutions, and luminescence was recorded using a GloMax 96 Microplate Luminometer (#E6521, Promega, Madison, United States). Adenylate concentrations were assessed by comparison to standard solutions of ATP disodium salt or ADP disodium salt.

**Determination of $P_i$ contents**

For the determination of free $P_i$, 20-60 mg of plant material were frozen in liquid nitrogen, ground with a bead mill, and resuspended in extraction buffer (10 mM Tris; 1 mM EDTA; 100 mM NaCl; 5 mM DTT; 1 mM phenylmethylsulfonyl fluoride; pH 8), using 10 µl per 1 mg fresh weight of plant material. 100 µl of the homogenized lysate were mixed with 900 µl of 1 % acetic acid and incubated at 42°C for 30 min. After pelleting cell debris (14,000 g, 10 min), 300 µl of the supernatant were mixed with 700 µl of freshly prepared assay solution (1.86 mM $(NH_4)_6MO_7O_{24}$; 80 mM ascorbic acid; 0.43 M $H_2SO_4$) and incubated at 42°C for 30 min. The absorbance of the solution was measured at 820 nm using a spectrophotometer. For calibration, $KH_2PO_4$ solutions were prepared in extraction buffer and processed in parallel.

**Protoplast isolation, transfection and transactivation assay**

Leaves from 4-6 weeks-old rosette plants were used for protoplast isolation. The lower epidermal cell layer was pulled off using Scotch Magic (3 M, Saint Paul, United States), while the upper leaf surface was fixed with ROTI Tape (Carl Roth, Karlsruhe, Germany). Peeled leaves were transferred into 5-10 ml of enzyme solution (0.4 M mannitol; 20 mM MES, pH 5.7; 10 mM $CaCl_2$; 20 mM KCl; 1 % (w/v) cellulase (#16419; SERVA Electrophoresis, Heidelberg, Germany); 0.25 % (w/v) macerozyme (#28302; SERVA Electrophoresis, Heidelberg, Germany); 0.1 % (w/v) bovine serum albumin), and incubated in the dark for 3 h. Subsequently, protoplasts were transferred to round-bottom cell culture tubes (#163160; Greiner Bio-One; Kremsmünster, Austria), pelleted by centrifugation at 100 *g*, 4°C, for 3 min, resuspended in 5 ml of pre-chilled W5 solution (154 mM NaCl; 125 mM $CaCl_2$; 5 mM KCl; 5 mM glucose; 2 mM MES, pH 5.7), and incubated on ice for 25 min. W5 solution was discarded from sedimented protoplasts, and protoplasts were resuspended to a final density of $3 \cdot 10^5$ cells/ml in pre-chilled MMG solution (0.4 M mannitol; 15 mM $MgCl_2$; 4 mM MES, pH 5.7).

For protoplast transfection, 2 µg of the reporter plasmid (pBT10-promoter::$LUC_{Firefly}$, containing promoter constructs of *MGD3*, *SPX1*, *GPT2*, or *SRG3*), 4 µg of the effector plasmids (pHBTL-p35S::GFP or pHBTL-p35S::PHR1), and 1 µg of the normalization vector pBT10-*pro35S::*$LUC_{Renilla}$ (Bäumler et al., 2019) were premixed. 400 µl of protoplast suspension was added to the plasmid mixture followed by addition of 440 µl of PEG solution (40 % (w/v) PEG4000 (#81242; Sigma-Aldrich, St. Louis, United States); 0.2 M mannitol, 0.1 M $CaCl_2$). After mixing by inversion for 4 min and incubation for 10 min, 1.76 ml of W5 solution were added, and protoplasts were pelleted by centrifugation for 1 min at 200 *g*, 4°C. Pellets were resuspended in 400 µl of W1 solution (0.5 M mannitol; 20 mM KCl; 4 mM MES, pH 5.7), and protoplasts were incubated in the dark at 19°C for 16 hours. For luminescence measurements, 200 µl of protoplast suspension was pelleted at 200 *g*, 4°C, 1 min, and lysed by the addition of 1x Beetle-Lysis juice (#10251, PJK, Kleinblittersdorf, Germany) and vortexing for 10 s. Cell debris was removed by centrifugation, and 20 µl of the supernatant were subjected to luminescence measurements after automated addition of 50 µl *Renilla*-Juice containing coelenterazine (Kit #102531, PJK), or Beetle-Juice containing D-luciferine and ATP (Kit #1025110, PJK) for the detection of *Renilla* or Firefly luciferase activity, respectively. Substrate addition and luminescence measurement was performed using the GloMax 96 Microplate Luminometer (#E6521, Promega, Madison, United States) with an integration time of 3 sec.
