## Supporting Table S4 for "PHR1 and PHL1 mediate rapid high-light responses and acclimation to high photosynthetic activity"

**Supporting Information Table S4** List of primer sequences used in this study. Non-annealing regions are indicated by shading.

| Primers | Sequence |
|---|---|
| **Genotyping primers** | |
| PHR1_F | 5’ AAA CGA AGT GAT TGG CAT GAA 3’ |
| PHR1_R | 5’ AAG CGG TGT CAA CTT CCT TTC 3’ |
| LBb1 | 5’ GCGTGGACCGCTTGCTGCAACT 3’ |
| PHL1_F | 5’ TCC CAC AAT CCA AAT TCA GAG 3’ |
| PHL1_R | 5’ CGG TTT ATA CCT TGC CGT TCT 3’ |
| LB3 | 5’ GAAT TTCA TAA CCA ATC TCG ATA CAC 3’ |
| NPTII_F | 5’ TAT GAC TGG GCA CAA CAG ACA 3’ |
| NPTII_R | 5’ TTCA GCA ATA TCA CGG GTA GC 3’ |
| TPT_F | 5’ GGT GAT ATG GAA CTT CGA TTG G 3’ |
| TPT_R | 5’ GCG GTA TCT CCA CCT TCA GC 3’ |
| *tpt-2*_F | 5’ GTA ACT TAC GAG TAA ACT GGC TAC 3’ |
| GPT2_F | 5’ CGG TTT CTC AAG TCG GAC CA 3’ |
| GPT2_R | 5’ TGC TTC GCC TGC TCA ATG AT 3’ |
| GK-LB | 5’ ATA TTG ACC ATC ATA CTC ATT GC 3’ |
| qSRG3_F | 5’ CGG AGG AAC TGA GAT GTA CC 3’ |
| SRG3_R | 5’ ATC CTT ACA GAC CTC TCT TGC CC 3’ |
| LBb1 | 5’ GCG TGG ACC GCT TGC TGC AAC T 3’ |
| **qRT-PCR primers** | |
| qPP2A_F | 5’ TAA CGT GGC CAA AAT GAT GC 3’ |
| qPP2A_R | 5’ GTT CTC CAC AAC CGC TTG GT 3’ |
| qSPX1_F | 5’ TCC AAG CAG AGT TAT CAG AGC AT 3’ |
| qSPX1_R | 5’ GGC GGC AAT GAA AAC ACA CT 3’ |
| qMGD3_F | 5’ ATC ACT AAG GCT GGT CCG GGT ACG 3’ |
| qMGD3_R | 5’ TGT CCA CAA CAT ACG GCA CGT TGC C 3’ |
| qSRG3_F | 5’ CGG AGG AAC TGA GAT GTA CC 3’ |
| qSRG3_R | 5’ CCG TAT GTC ATC AGA GAG AGC 3’ |
| qVPE1_F | 5’ CCT TAC TAA CGG GTT ACA TAT CG 3’ |
| qVPE1_R | 5’ GGC CTA GAC TCA GCT ATC TTC 3’ |

| | |
|---|---|
| qPS2_F | 5’ CTT GCC CTC CTA ACA TGT GC 3’ |
| qPS2_R | 5’ CTT GGA CAG TAA TCG CCA GC 3’ |
| qbZIP63_F | 5’ CGT TGA ATC GCA GTG CTT CC 3’ |
| qbZIP63_R | 5’ GGA GAC GGA AAC ACC ACA CG 3’ |
| qLDOX_F | 5’ TGA GCT AGC ACT CGG TGT GG 3’ |
| qLDOX_R | 5’ AAA GCT GCA AAC CCG GAA CC 3’ |
| qDFR_F | 5’ GGG TTT CAT CGG TTC ATG G 3’ |
| qDFR_R | 5’ AGT AGC GTC TTG GCG TTT GG 3’ |
| qPAP1_F | 5’ CTG GTC GGA CCG CAA ATG A 3’ |
| qPAP1_R | 5’ GGT GTT GTA GGA ATG GGC GT 3’ |
| qPAP2_F | 5’ GCC ACA ATA ACC CCC TAT TCC 3’ |
| qPAP2_R | 5’ CTC AAC CCT TTG GAC GAA CC 3’ |
| qMYB111_F | 5’ GAC CGA GAA GCA ATG GGA AG 3’ |
| qMYB111_R | 5’ CTT CCT CGG CTG TCC ATC TC 3’ |
| qHSP70_F | 5’ TGGGAATCAACTGGCTGAGG 3’ |
| qHSP70_R | 5’ TATCAGGCCCAGCTCCTTGG 3’ |
| qHSP18.2_F | 5’ AGC GGA GAG AGG AGC AAG GA 3’ |
| qHSP18.2_R | 5’ CGG AAC CAC AAC CGT AAG CA 3’ |
| **Primers used for cloning** | |
| proGPT2_F | 5’ ACC TGG ATC CGA ATG AAA ATG ACA AAC GAT ACA TTG 3’ |
| proGPT2_R | 5’ ATT GCC ATG GTG TGC TTT TTT ATG GCT AAT TGA TGA 3’ |
| proSRG3_F | 5’ ACC TGG ATC CCG CAG GTT GTC GAT ACA AAA G 3’ |
| proSRG3_R | 5’ ATT GCC ATG GAT TTC TAT TTT TAG AAA GAA AAA AGG GC 3’ |
| proSPX1_F | 5’ ACC TGA ATT CGT CGG TTC GGT TTG GTT CTG 3’ |
| proSPX1_R | 5’ ATT GCC ATG GAG CTC TTT TAT TTT CTG GGA AAC TTA A 3’ |
| proSPX1_mut_F | 5’ CAA GAA TAT TCC ATC GAA TCC AAC 3’ |
| proSPX1_mut_R | 5’ GTT GGA TTC GAT GGA ATA TTC TTG 3’ |
| attB1-HA_FW | 5’ AAA AAG CAG GCT TAA TGG GAT CCT ACC CAT ACG 3’ |
| PHR1-HA_R | 5’ CGA GCC TCT CCA GCA GAT CCA GCG TAA TCT GGA ACG TCG TAT G 3’ |
| HA-PHR1_F | 5’ GAT CTG CTG GAG AGG CTC GTC CAG TTC ATA G 3’ |
| attB2-PHR1_R | 5’ AGA AAG CTG GGT AGC CAG GTT TAC TAT TTA CTC ATA 3’ |
| attB1_F | 5’ GGG GAC AAG TTT GTA CAA AAA AGC AGG CT 3’ |
| attB2_R | 5’ GGG GAC CAC TTT GTA CAA GAA AGC TGG GT 3’ |
