## Supporting Figures for "PHR1 and PHL1 mediate rapid high-light responses and acclimation to high photosynthetic activity"

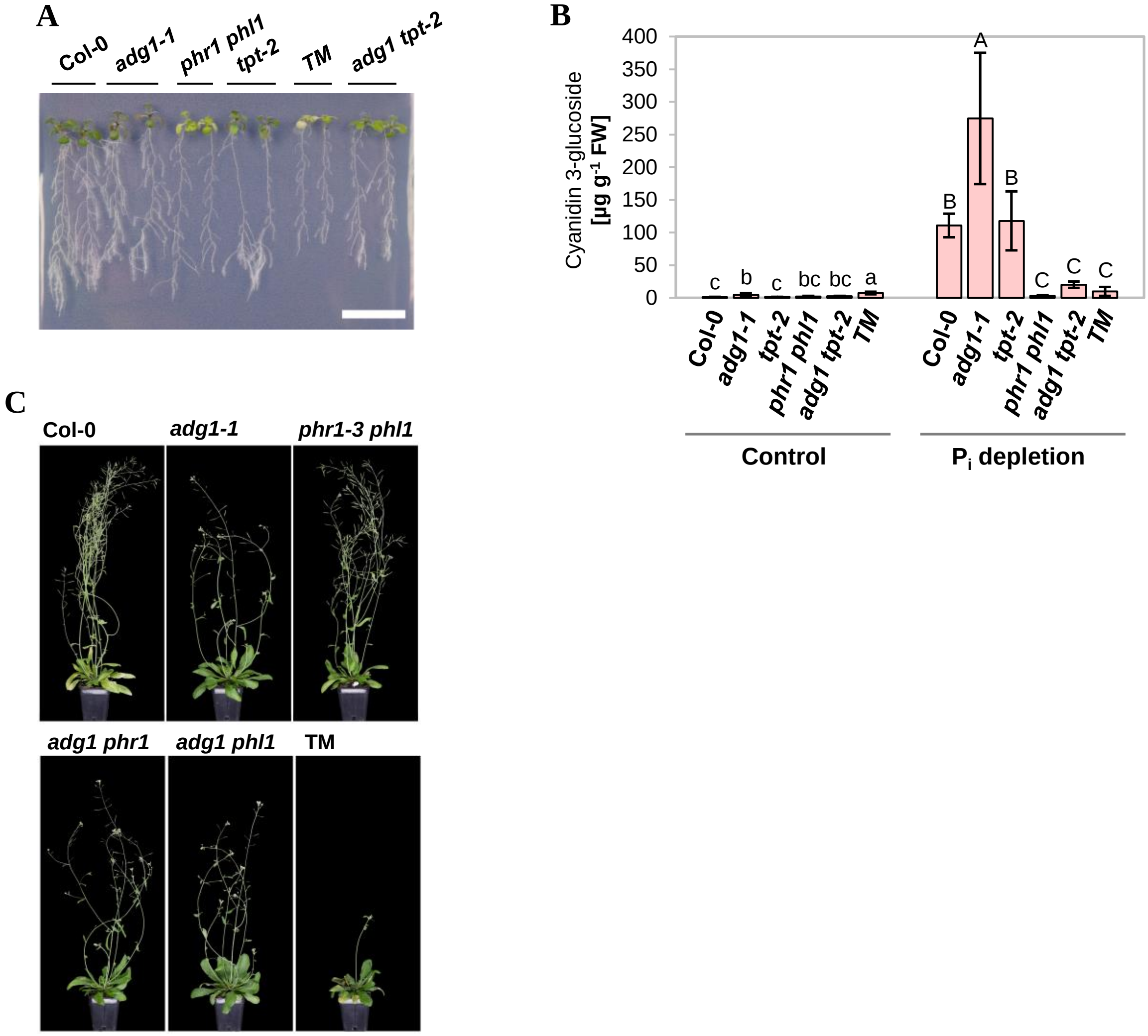


**Supporting Information Fig. S1** Phenotypes of *adg1-1 phr1-3 phl1* (**TM**). **A,** Seedling phenotypes under $P_i$ deficient growth conditions. Seedlings of WT (Col-0), *adg1-1*, *tpt-2*, *phr1-3 phl1*, *adg1-1 tpt-2* and TM genotype were grown for 7 days on rich medium (½ MS) including 0.5 % sucrose before transfer to media with 0.5 % sucrose and either 2.5 mM (**Control**) or 0 mM (**$P_i$ depletion**) $KH_2PO_4$ added. Growth was continued for 8 days. Bar, 2 cm. **B**, Anthocyanin (cyanidin 3-glucoside) contents of seedling shoots. Seedlings were grown and treated as described in A, but transfer to $P_i$-deferring media was conducted after 5 days of growth and treatment was sustained for 7 days. Pigment levels are depicted relative to shoot fresh weights. Bars represent means ± standard deviations; $n$ = 5-6 pools of seedling shoots from 3 independent experiments. One-way ANOVA with Tukey HSD follow-up test and Bonferroni alpha correction for contrasts; $P$ <0.05. **C**, Inflorescences of WT (Col-0), *adg1-1*, *phr1-3 phl1*, and derived genotypes showing delay of the transition to flowering in the TM. Plants were grown under a 16-h light regiment at 23°C and a light intensity of 100±10 µmol $m^{-2}$ $s^{-1}$. Pictures were taken of representative plants after 42 days of growth. Backgrounds were manually removed for better visualization.

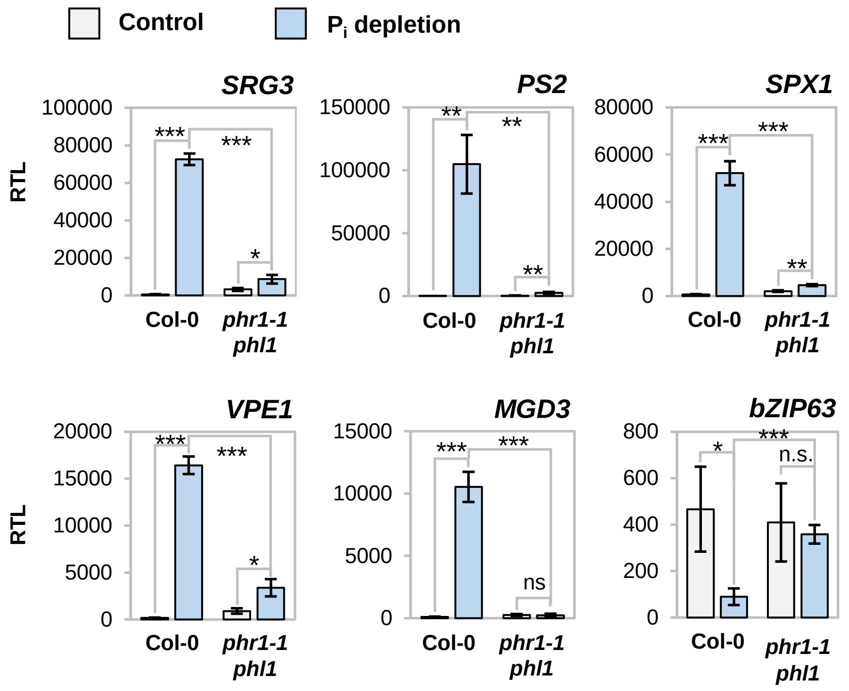


**Supporting Information Fig. S2** Expression of selected $P_i$-starvation responsive genes depends on PHR1/PHL1 under $P_i$ depletion. qRT-PCR analysis of *SRG3*, *PS2*, *SPX1*, *VPE1*, *MGD3*, and *bZIP63* transcript levels in seedlings of WT (Col-0) and *phr1-1 phl1* mutant genotype. Seedlings were grown on rich medium for 7 days before transfer to media with either 2.5 mM (Control, grey bars) or 0 mM ($P_i$ depletion, blue bars) $KH_2PO_4$ added. Shoot material was harvested after additional 8 days of growth 9 h after onset of the 16-h photoperiod. Transcript levels were calculated relative to the transcript levels of *PP2A* as $1000 \cdot 2^{-\Delta CT}$. Bars show means ± standard deviations. $n$ = 3 independent experiments. One-way ANOVA with Tukey HSD follow-up test and Bonferroni alpha correction for contrasts; ***$P$ <0.001, **$P$ <0.01, *$P$ <0.05, n.s., not significant.

**A**

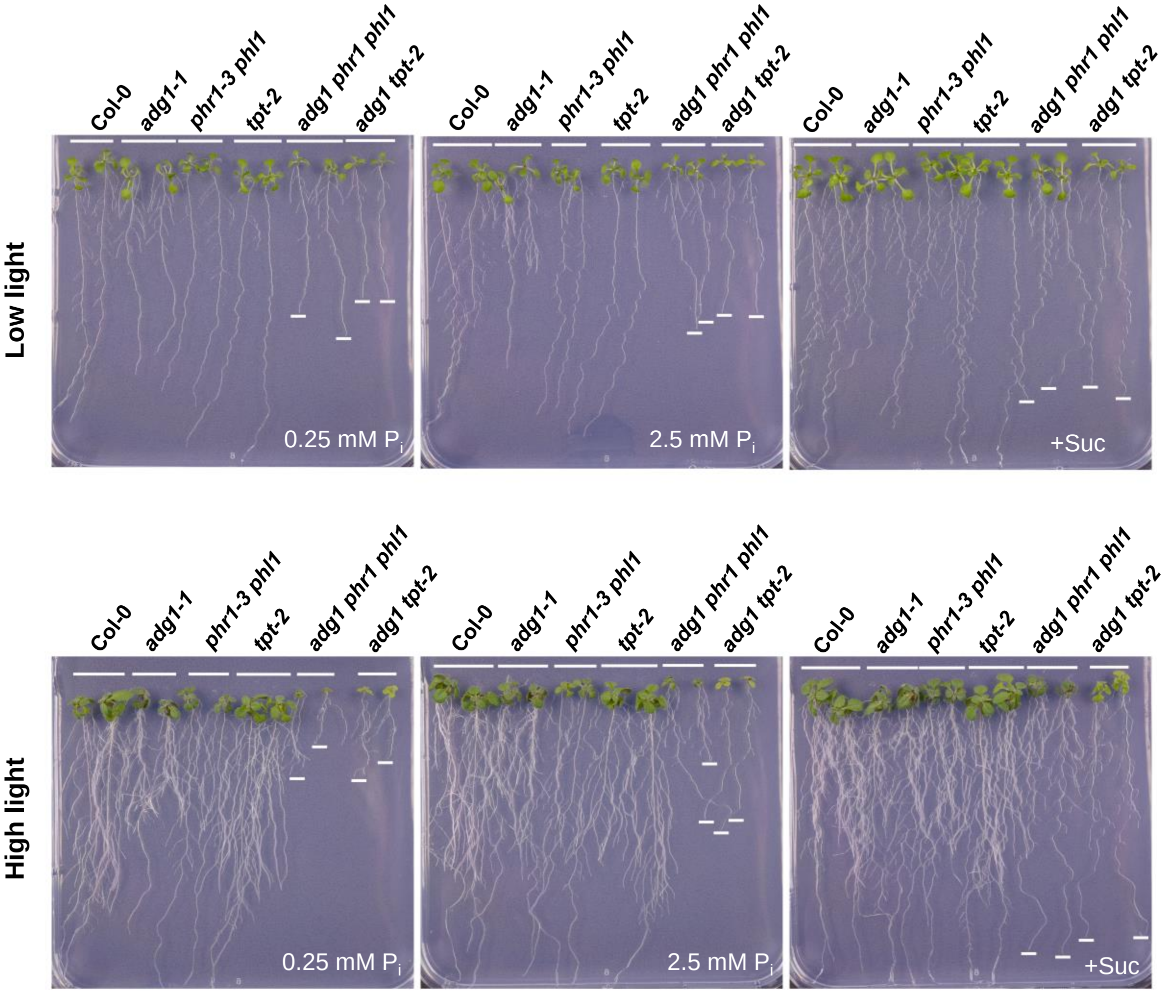


**Supporting Information Fig. S3** Root growth of *adg1 phr1 phl1* and *adg1 tpt-2* responds to exogenously applied sucrose. Phenotypes of Col-0, *adg1-1*, *phr1-3 phl1*, *tpt-2* and derived mutant lines. Seedlings were grown as described in Figure 1F. White bars indicate apical ends of the primary roots of *adg1 phr1 phl1* and *adg1 tpt-2* lines. The experiment was performed 3 times with each 8 seedlings per condition. Representative pictures are shown.

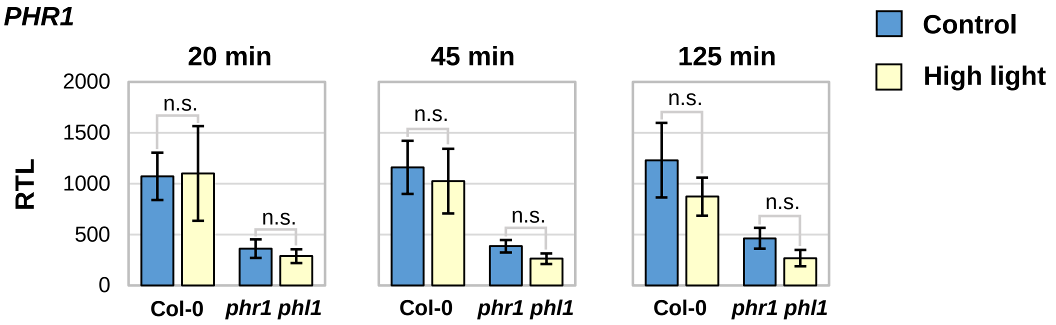


**Supporting Information Fig. S4** *PHR1* transcript levels are not affected by short-time high-light exposure. qRT-PCR analysis of *PHR1* transcripts in rosette leaves from WT (Col-0) and *phr1-1 phl1* mutants. Plants were grown and treated as described in Figure 2B. Transcripts were calculated relative to *PP2A* as $1000 \cdot 2^{-\Delta CT}$. Residual *PHR1* expression seen in *phr1-1 phl1* double mutants corresponds to missense transcript containing an early stop codon (Bustos et al., 2010). Bars represent means ± standard deviations; $n$ = 3 independent experiments. Statistical analyses were performed using Student's $t$ test with 2-tailed distribution; n.s., not significant.

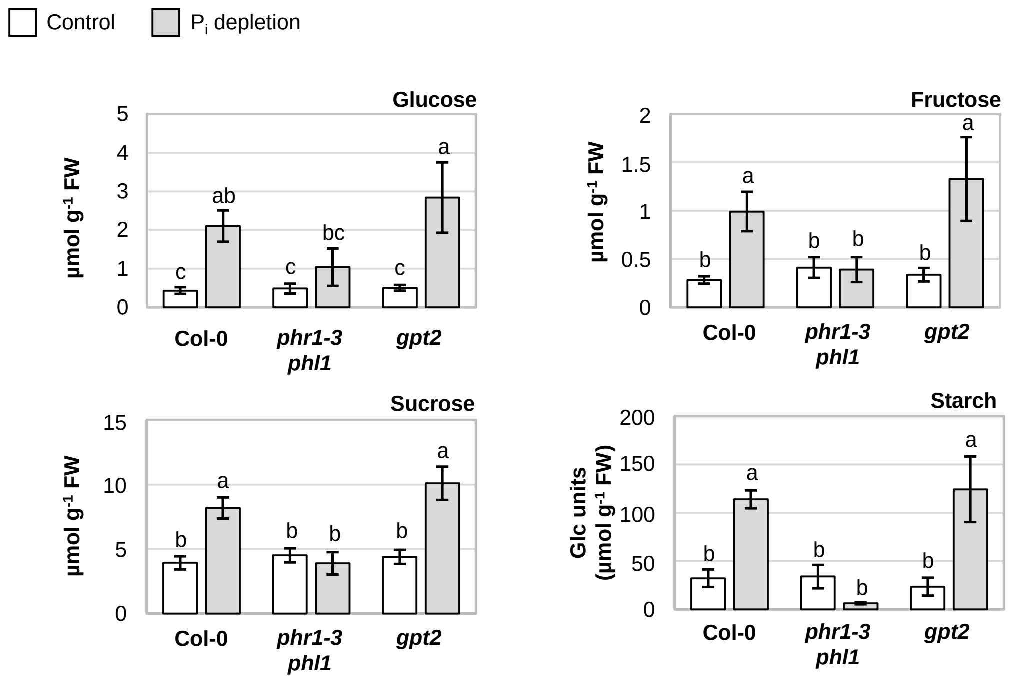


**Supporting Information Fig. S5** Changes in sugar and starch contents under $P_i$ depletion. Seedlings of WT (Col-0), *phr1-3 phl1* and *gpt2* mutant genotype were grown for 10 days on rich medium before transfer to media containing either 2.5 mM (Control, white bars) or 0 mM ($P_i$ depletion, grey bars) $KH_2PO_4$. Seedling shoots were harvested 10.25 h after onset of the 16-h photoperiod. Contents of glucose, fructose, sucrose and starch were determined after additional 7 days of growth. Bars represent means ± standard deviations; $n$ = 3 independent experiments; 2-factor ANOVA with Tukey HSD post-hoc test; $P < 0.05$.

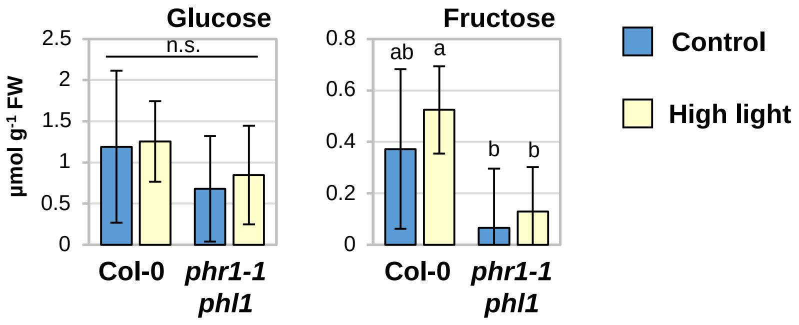


**Supporting Information Fig. S6** Monosaccheride levels after 20 min of high light. Plants were grown and treated as described in Figure 2B. *n* = 8 plants from 4 independent experiments; bars show means ± standard deviations; 2-factor ANOVA with Tukey HSD post-hoc test; $P < 0.05$, n. s. not significant.

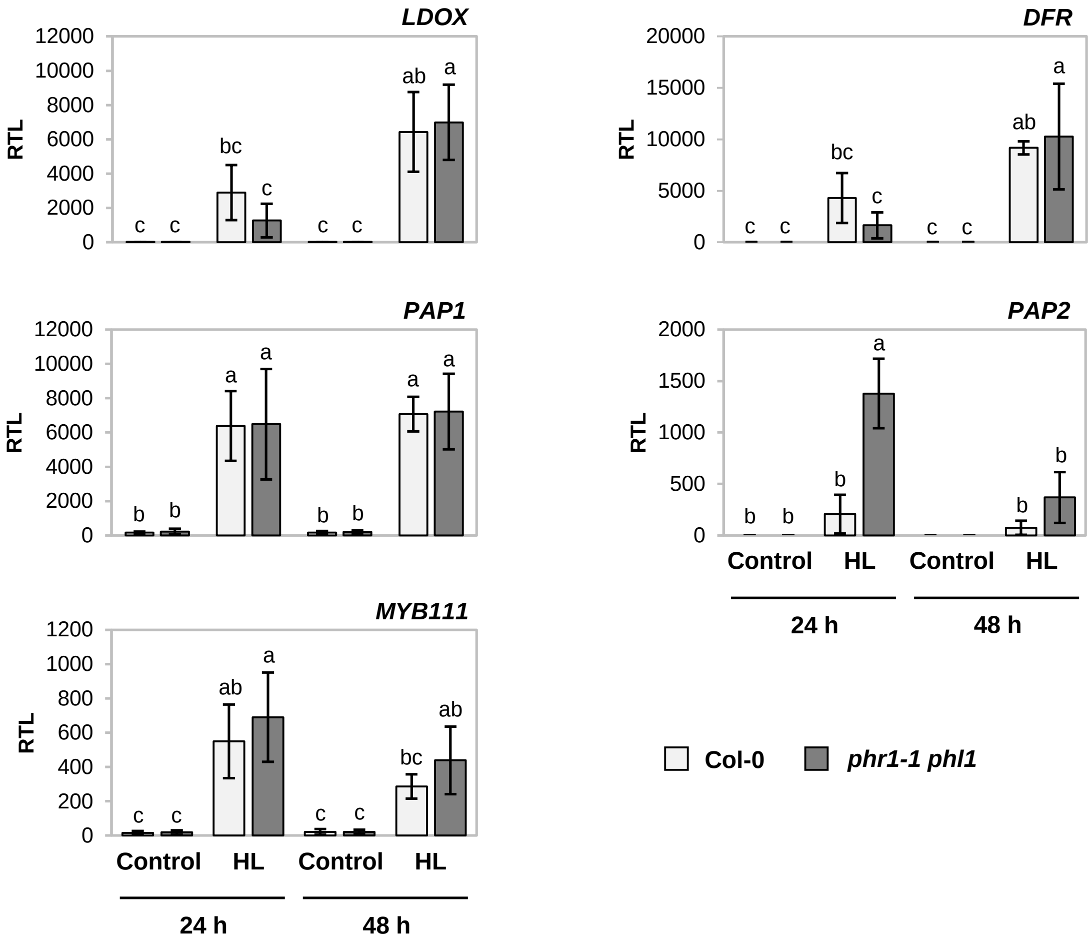


**Supporting Information Fig. S7** Anthocyanin biosynthetic and regulatory gene expression upon high light in WT and *phr1-1 phl1*. WT (Col-0) and *phr1-1 phl1* mutants were grown for 39 days as described for Fig. 2B before light intensity was shifted to 450±30 µmol $m^{-2}$ $s^{-1}$ (high light) at 4 h after onset of the photoperiod. Control plants were kept under growth light conditions (70±5 µmol $m^{-2}$ $s^{-1}$). Material was harvested after 24 and 48 h. Transcript levels of *LDOX*, *DFR*, *PAP1*, *PAP2*, and *MYB111* were calculated relative to *PP2A* as $1000 \cdot 2^{-\Delta CT}$. Bars show means ± standard deviations; $n$ = 3 independent experiments; 2-factor ANOVA with Tukey HSD post-hoc test; $P < 0.05$.

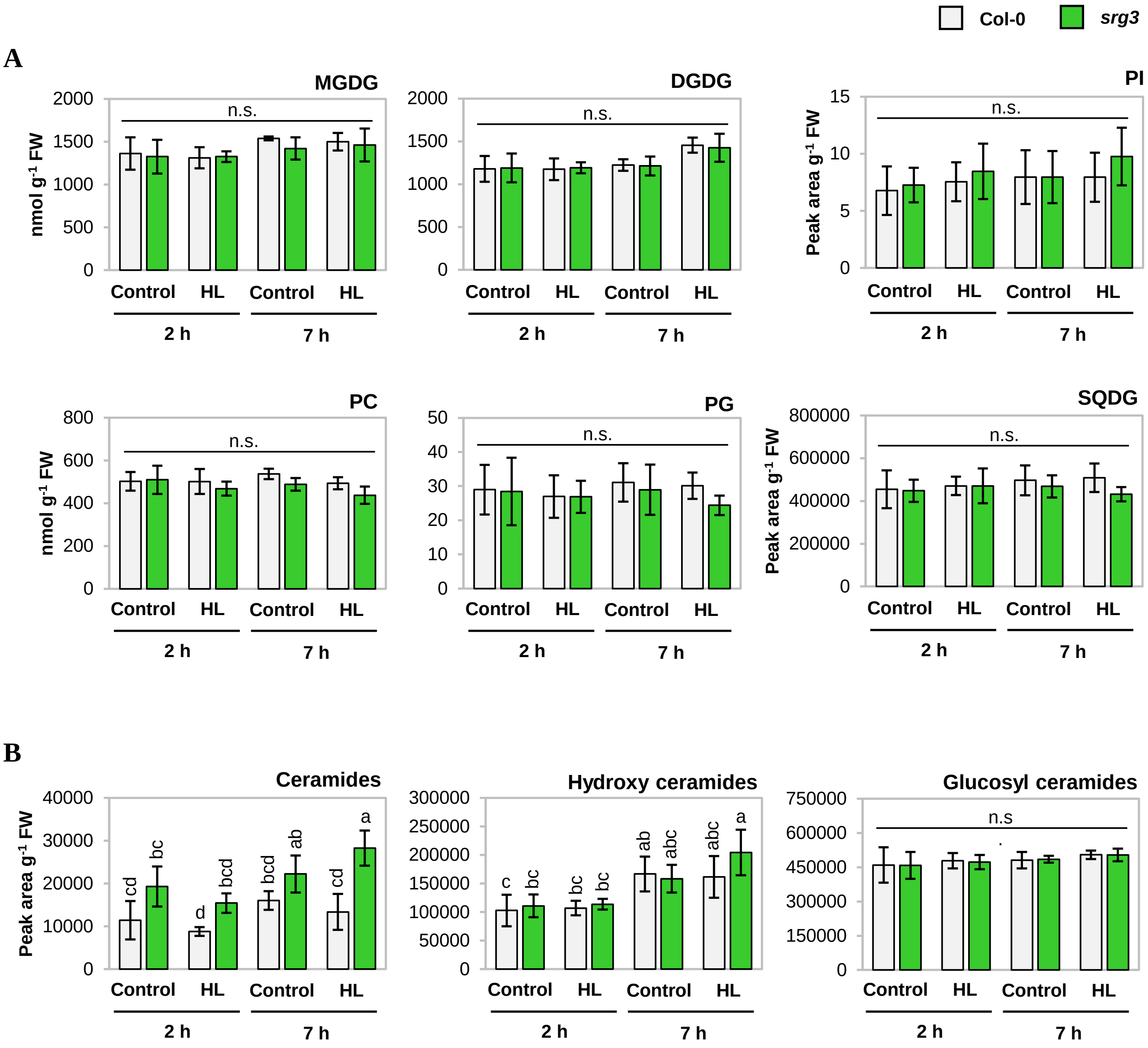


**Supporting Information Fig. S8** Total levels of lipid classes in WT (Col-0) and *srg3* mutants upon shift to high light (**HL**). Plants were grown and treated as described for Fig. 2B. Rosette leaves (2-3 per sample) were harvested after 2 or 7 h of treatment. Contents were calculated relative to fresh weights (FW). Bars represent means ± standard deviations; $n = 4$ independent experiments; 2-factor ANOVA with Tukey HSD post-hoc test; $P < 0.05$; n.s., not significant. **A**, MGDG, Monogalactosyl diacylglycerol; DGDG, Digalactosyl diacylglycerol; PI, Phosphatidylinositol; PC, Phosphatidylcholine; PG, Phosphatidylglycerol; SQDG, Sulfoquinovosyl diacylglycerol. **B**, Levels of ceramides and of the ceramide derivatives hydroxy ceramides and glucosyl ceramides.

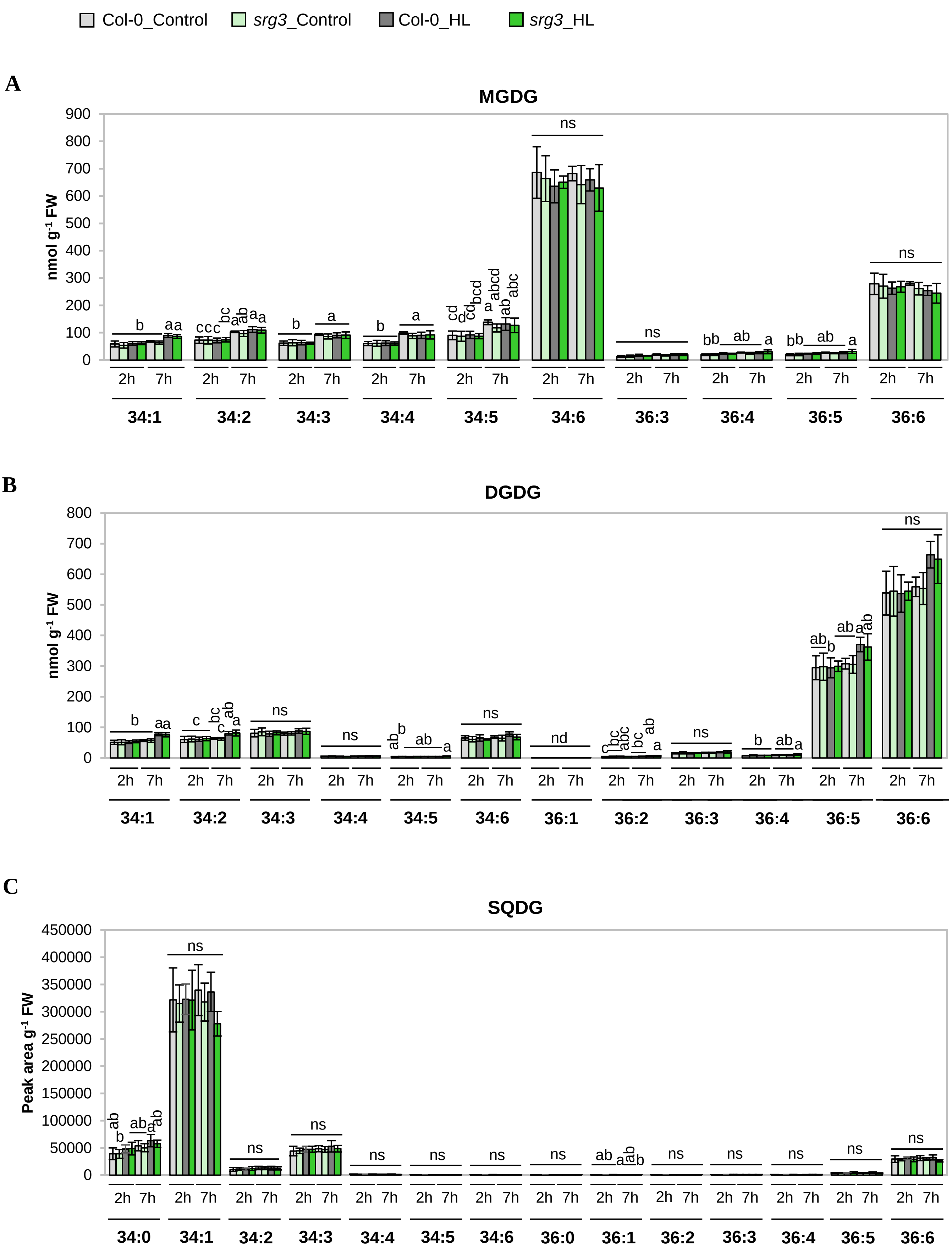


**Supporting Information Fig. S9** Contents of glycosylglycerol lipid species in WT (Col-0) and *srg3* mutants upon shift to high light (**HL**) relative to fresh weight (FW). Plants were grown an treated as described for Supplemental Figure 8. Bars represent means ± standard deviations; $n = 4$ independent experiments; 2-factor ANOVA with Tukey HSD post-hoc test; $P < 0.05$; n.s., not significant. MGDG, Monogalactosyl diacylglycerol; DGDG, Monogalactosyl diacylglycerol; SQDG, Sulfoquinovosyl diacylglycerol.

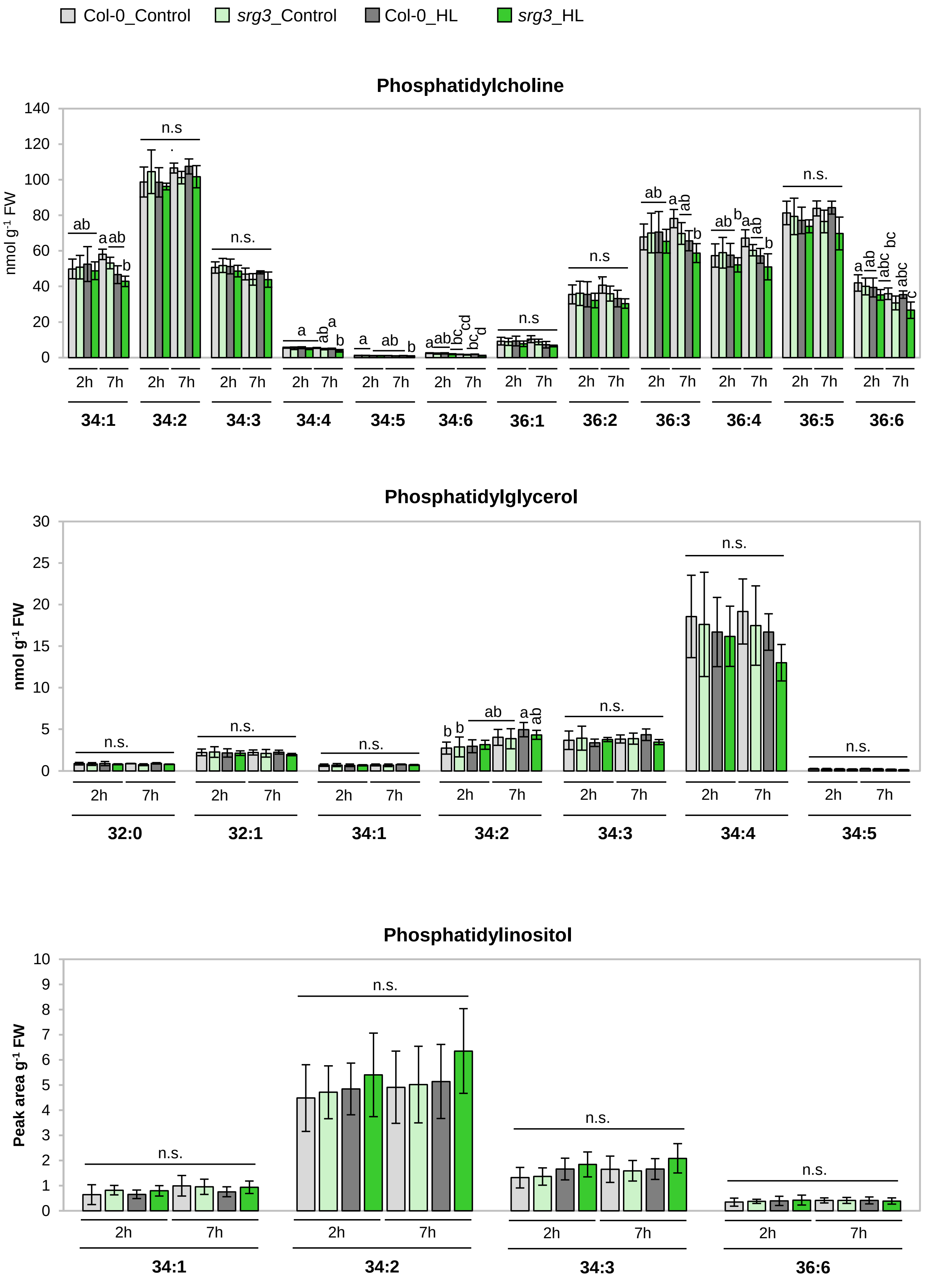


**Supporting Information Fig. S10** Contents of glycerophospholipid species in WT (Col-0) and *srg3* mutants upon shift to high light (**HL**) relative to fresh weight (FW). Plants were grown an treated as described for Supplemental Figure 8. Bars represent means ± standard deviations; $n = 4$ independent experiments; 2-factor ANOVA with Tukey HSD post-hoc test; $P < 0.05$; n.s., not significant.

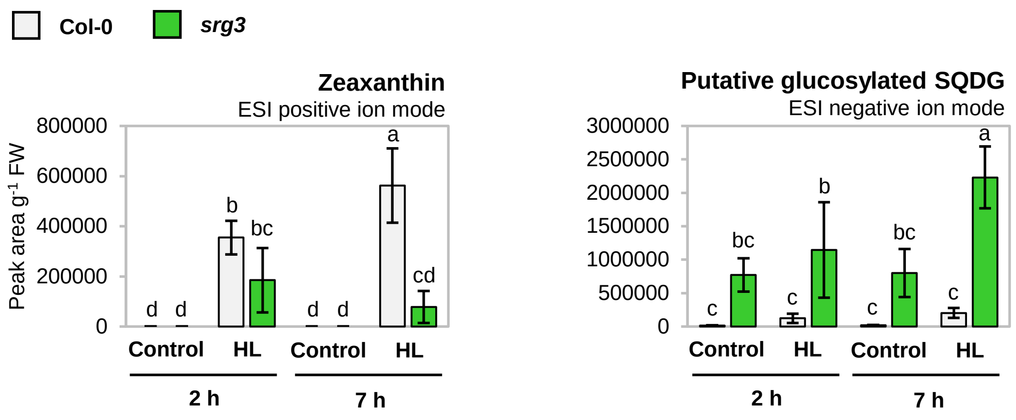


**Supporting Information Fig. S11** Levels of zeaxanthin and putative glycosylated SQDG determined in opposite ESI ion mode compared to Fig. 5C. Plants of WT (Col-0) and *srg3* mutant genotype were grown an treated as described for Supplemental Figure 8. **HL**, High light. Bars represent means ± standard deviations; $n = 4$ independent experiments; 2-factor ANOVA with Tukey HSD post-hoc test; $P < 0.05$.

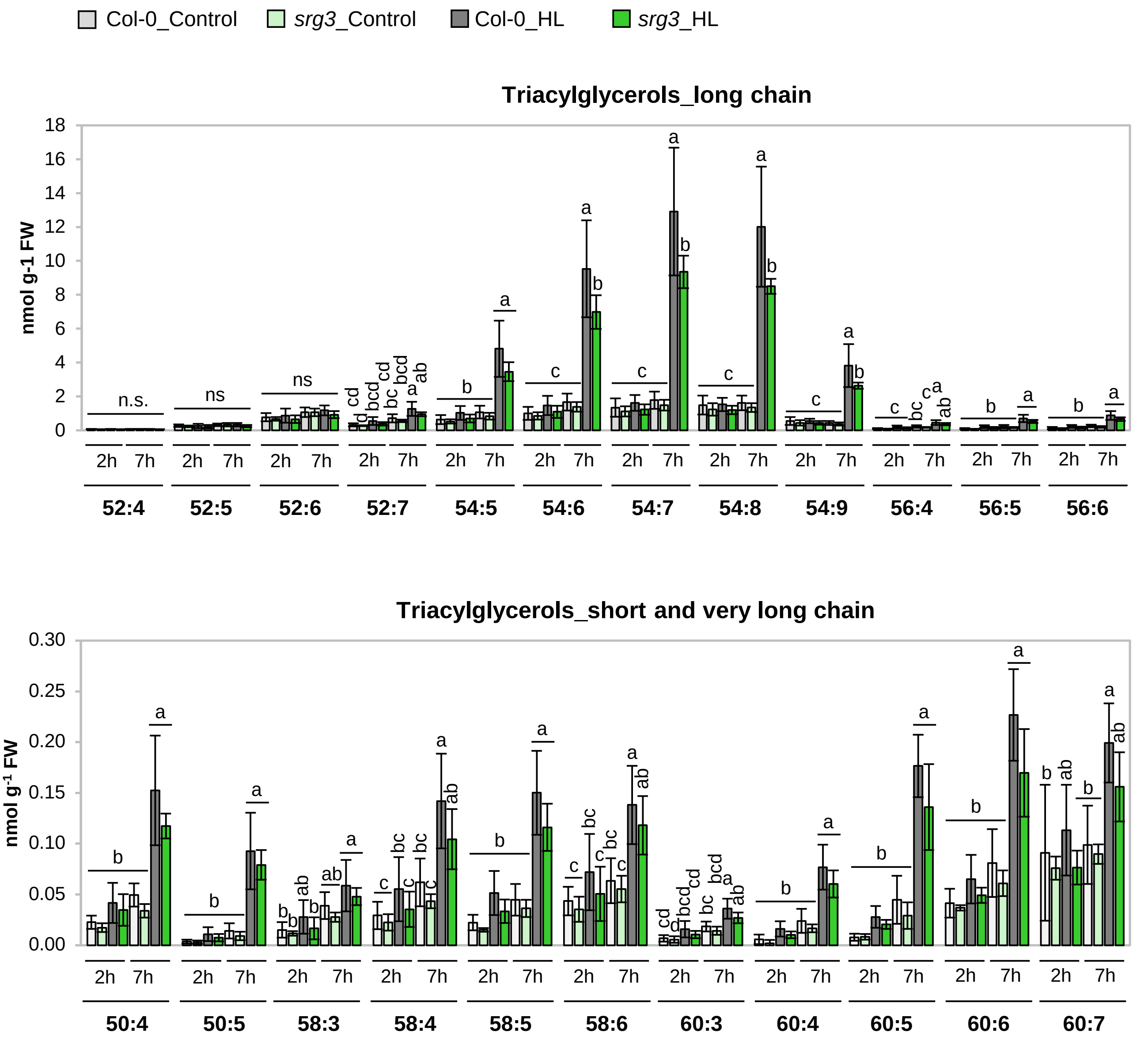


**Supporting Information Fig. S12** Levels of triacylglycerol species in WT (Col-0) and *srg3* mutants upon shift to high light (**HL**). Plants were grown and treated as described for Supplemental Figure 8. Bars represent means ± standard deviations; $n = 4$ independent experiments; 2-factor ANOVA with Tukey HSD post-hoc test; $P < 0.05$.

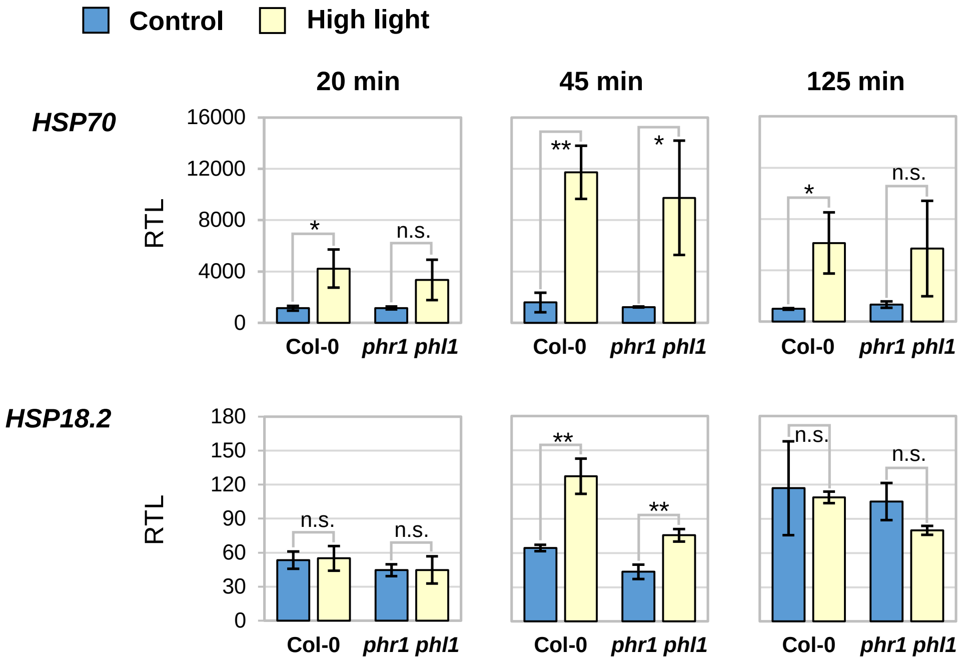


**Supporting Information Fig. S13** Levels of high-temperature responsive transcripts *HSP70* and *HSP18.2* in WT (Col-0) and *phr1-1 phl1* mutant plants upon high light treatment. Plants were grown and treated as described in Fig. 2B. Rosette leaves were sampled after 20, 45, and 125 min of treatment. Transcript levels were calculated relative to *PP2A* as $1000 \cdot 2^{-\Delta CT}$. Bars represent means ± standard deviations, $n = 3$ independent experiments. Statistical analyses were performed using Student's $t$ test with 2-tailed distribution, unpaired with unequal variance. Significant differences are indicated as $^{**}P < 0.01$, $^{*}P < 0.05$, n.s., not significant.
